# Recent effective population size of the snowy owl (*Bubo scandiacus*) tracks historical climate variation

**DOI:** 10.64898/2026.09.30.752484

**Authors:** E.L.G. Enevoldsen, D. Aguilar-Gómez, K.-O. Jacobsen, R. Solheim, I.J Øien, T. Aarvak, S.N.K. Hoff, O.S. Kersten, K.L Wiebe, J.F. Therrien, I.E. Menyushina, R. Nielsen, S. Boessenkool, H.T. Baalsrud

**Affiliations:** Department of Biosciences, University of Oslo, Oslo, Norway; Department of Animal and Aquacultural Sciences, Norwegian University of Life Sciences, Ås, Norway; Department of Ecology and Evolutionary Biology, University of California Los Angeles, CA, USA; Norwegian Institute for Nature Research, Fram Centre, Tromsø, Norway; Natural History Museum, University of Agder, Kristiansand, Norway; Department of Biology, University of Saskatchewan, Saskatoon, Canada; Acopian Center for Conservation Learning, Hawk Mountain Sanctuary, Orwigsburg, PA, United States; BirdLife Norway, Trondheim, Norway; Department of Integrative Biology, University of California Berkeley, California, USA

**Keywords:** snowy owl, conservation genomics, genetic diversity, population structure, panmixia, centromeres, Arctic, circumpolar

## Abstract

Arctic ecosystems are undergoing rapid climate warming, providing an opportunity to examine molecular evolution under environmental stress. Here, we used whole-genome resequencing to assess population structure, genetic diversity, and recent demographic history in the snowy owl (*Bubo scandiacus*), a highly mobile Arctic apex predator classified as Vulnerable by the IUCN. We analysed genome-wide data from individuals sampled across the species’ circumpolar range using a chromosome-level reference genome. Population genomic analyses revealed an absence of detectable population structure, indicating extensive gene flow and near-panmixia across continents. Genome-wide nucleotide diversity was low (π = 2.47 × 10^−4^), while runs of homozygosity were dominated by short tracts, indicating limited standing genetic variation despite little evidence for recent close inbreeding. Our demographic inference revealed a sustained decline in effective population size over the past ~1,600 years, a pronounced reduction coinciding with the Medieval Warm Period followed by a partial recovery during the subsequent cooler interval. This direct, measurable effect of past climate warming on effective population size raises concerns for the long-term survival of the snowy owl. Our results demonstrate that high connectivity and low levels of recent inbreeding have not necessarily buffered this species against climate-driven demographic erosion of genetic diversity. This study highlights the value of using whole-genome population genomics to inform genomic monitoring and conservation decision-making in rapidly changing environments.

## 1. Introduction

Arctic ecosystems are undergoing rapid and profound climate change, with significant but often unpredictable consequences for wildlife. One iconic species that shows declining population trends is the snowy owl (*Bubo scandiacus*). The IUCN recently reclassified the snowy owl from “Least Concern” to “Vulnerable” [1] following rapid population declines, and current estimates suggest a global population of 14,000–28,000 individuals [2]. Occupying the very northern edges of our planet, the snowy owl is a tundra-obligate species with nowhere to escape to under a warming climate. The species displays an extraordinary degree of mobility, with banding and satellite telemetry showing that single individuals can breed thousands of kilometres apart in different years [3–5]. Snowy owls are long-lived for a bird their size, with banding records showing wild individuals surviving more than 20 years [6], and annual adult survival estimated at 85-92% from satellite telemetry [2,7]. Breeding is tightly coupled to prey availability: pairs typically initiate nests only once lemming density exceeds a threshold [3,8].

Previous genetic work on North American, Fennoscandian and Siberian snowy owls did not find population structure, suggesting a high degree of gene flow across the Arctic [9], yet this was based on two mitochondrial genes which may lack sufficient power to detect barriers to gene flow in such a highly mobile species. Lack of population structure was also observed in North American snowy owls using RADseq [10]. Notably, in North America, regular southward migration to more temperate prairie regions [11] as well as overwintering in the Arctic [3,12] have been observed. By contrast, snowy owls nesting in Fennoscandia appear to be more nomadic, shifting locations across years in response to prey availability, but remaining in the Arctic throughout winter [13,14].

Recent technological developments now permit population-scale, whole-genome analyses that substantially improve the resolution and accuracy of population genomic and demographic inference [15]. This is especially consequential for highly mobile species, where fine-scale barriers to gene flow can be subtle and easily overlooked using a limited number of markers; resolving such barriers typically requires genome-wide panels comprising thousands of markers, rather than the handful of microsatellite loci or short mitochondrial fragments used in earlier studies [16–18]. High-resolution population genomic data allows us to better reconstruct the demographic history, population structure, and estimate effective population size (Ne) of snowy owls, which is central to assessing extinction risk and long-term viability [19]. Genome-wide diversity further provides insights into its evolutionary potential, i.e. its capacity to adapt to ongoing environmental change [20,21], and is increasingly urgent given the accelerating pace of Arctic warming, which has proceeded at nearly four times the global average rate over the past four decades [22].

Here, we use whole-genome resequencing data from 68 snowy owls sampled across their global distribution to assess range-wide genetic connectivity, recent demographic history, genome-wide genetic diversity and inbreeding. Moreover, we assessed whether any genomic regions show elevated differentiation among sampling localities. By combining this extensive geographic and genomic coverage with our recently published chromosome-level reference genome [23], we provide the most comprehensive population genomic assessment of the species to date. Our study provides critical new insights into the demographic history and resilience for one of the Arctic’s most wide-ranging predators.

## 2. Material and Methods

### (a) Sample information

Tissue samples from 68 snowy owls were collected from Norway and Sweden (hereby referred to as Fennoscandia), Canada, USA, Russia, and Greenland between 1996 and 2021 (Fig. 1). They consisted of either blood (n = 10), muscle tissue (n = 27), or feathers (n = 31). For detailed sample information, including sample ID, sex, life stage, location, and sampling year, see electronic supplementary materials, Table S1. A more detailed description of sampling procedures and molecular methods is provided in the electronic supplementary materials, Text S1.

**Figure 1.**
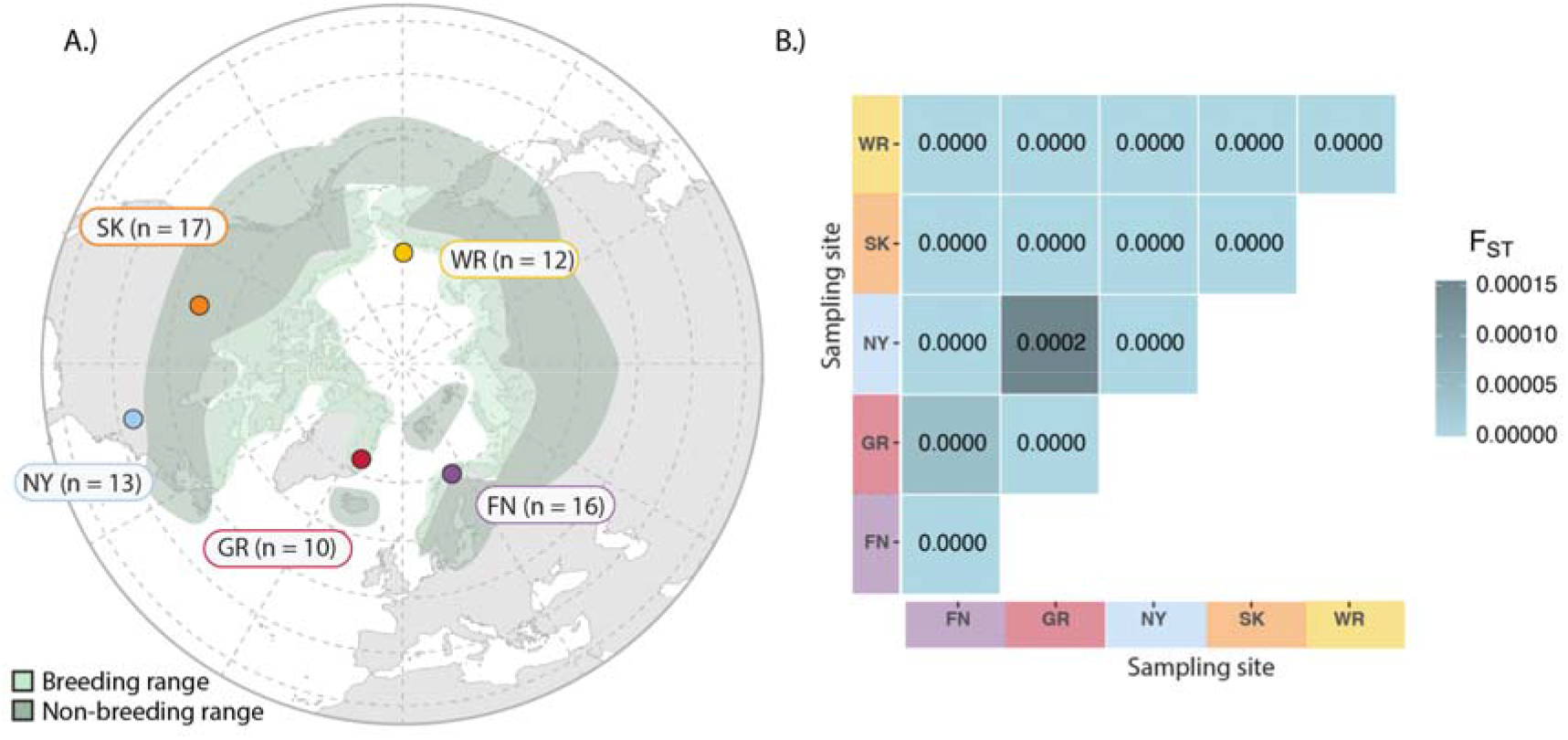
Snowy owl sampling locations and genetic structure. **A.** Sampling locations included in the population genomic analyses, with sample sizes shown next to each point. Shading indicates the breeding and non-breeding range of the snowy owl. Location abbreviations are FN, Fennoscandia; GR, Greenland; NY, New York; SK, Saskatchewan; and WR, Wrangel Island. **B.** Pairwise genetic differentiation among sampling locations, estimated as F_ST_ from autosomal pixy analyses of 50 curated individuals (FN = 13, GR = 8, NY = 13, SK = 11, WR = 5). Negative pairwise means are shown as zero.

### (b) Mapping, variant calling, and general filtering

Sequencing reads were processed using standardized population genomic pipelines. Detailed bioinformatic procedures are provided in the electronic supplementary materials, Text S1. Briefly, reads were trimmed with Trimmomatic v0.39 [24] and aligned to the chromosome-level snowy owl reference genome [23] using BWA-MEM v0.7.17 [25]. Alignment statistics were generated with Qualimap v2.3 [26] and summarized using MultiQC v1.14 [27].

Variants were called following GATK Best Practices using HaplotypeCaller and joint genotyping with GenotypeGVCFs (GATK v4.4; [28,29]). SNPs were hard-filtered using GATK-recommended thresholds (see electronic supplementary materials, Text S1. Following variant calling, autosomes and sex chromosomes were processed separately. Genotypes with depth <5x or genotype quality (GQ) <20 were set to missing, and site-level summary statistics were recalculated. For the MAF-filtered SNP dataset, autosomal sites were retained if they had a minor allele frequency ≥0.02 and missingness ≤20%. A separate dataset without a MAF threshold was prepared for all pixy analyses, as described below.

Quality-control analyses identified two samples (GRL04 and SKW02) that appeared to be contaminated (electronic supplementary materials, Text S1), and these individuals were excluded from all downstream analyses. The MAF-filtered autosomal SNP dataset contained 1,324,912 SNPs. Further sample curation was applied separately for PCA, pixy and demographic inference; analysis-specific inclusion is reported in Table S2.

### (c) Population structure

Population structure was assessed using multiple complementary approaches based on genome-wide SNP data. Principal component analysis (PCA) was performed using PLINK/PLINK2 [30,31] on a curated set of 50 individuals after filtering for missingness, contamination and relatedness (see below), retaining one representative from the closely related Fennoscandia group (electronic supplementary materials, Text S2 and Table S2). Prior to analysis, SNPs were pruned for linkage disequilibrium after retaining sites with minor allele count ≥3, using a sliding window of 50 SNPs, a step size of 10 SNPs, and an *r*² threshold of 0.2. Eigenvectors and eigenvalues were calculated using the PLINK --pca function and visualized in R v4.5.1 [32].

To evaluate whether variation in sequencing depth among individuals influenced population structure inference, a second PCA was performed using pseudohaploid genotypes generated with ANGSD v0.941 [33]. Pseudohaploid genotypes were obtained by randomly sampling a single allele at each site across individuals. The resulting pseudohaploid genotype matrix was analysed in R: biallelic polymorphic sites were retained, genotypes were coded as major and minor alleles, sites were mean-centred, missing values were set to zero after centring, and principal components were obtained by eigendecomposition of the sample covariance matrix. For the final autosomal pseudohaploid PCA, we used the curated PCA-filtered sample set and retained sites with minor allele count ≥ 3. As diagnostic analyses, we also performed PCA separately on Z-linked and W-linked pseudohaploid genotypes generated with ANGSD. Sites were filtered as for the autosomal ANGSD PCA. The W-chromosome PCA was restricted to genetically female individuals because males are not expected to carry W-linked sequences. Sex-linked PCA results were used only to assess potential sex-linked structure and technical artefacts, and were not treated as primary evidence for population structure.

### (d) Relatedness and genetic differentiation

Pairwise genetic similarity among individuals was estimated using the KING kinship statistic implemented in PLINK v2.0 [34]. Kinship coefficients were calculated from the LD-pruned autosomal SNP dataset and visualized as a heatmap of pairwise genetic similarity in R. Genetic differentiation and absolute divergence among sampling localities were quantified using Weir and Cockerham F_ST_ and dXY estimates from pixy v2.0.0 [35] in non-overlapping 50 kb autosomal windows, using the curated 50-individual all-sites dataset without a SNP MAF threshold. F_ST_ summaries used windows containing at least 10 SNPs and were weighted by the number of SNPs; dXY summaries were weighted by the number of callable sites. Negative F_ST_ estimates were retained in summaries and interpreted as no evidence of differentiation. Pairwise mean F_ST_ values were visualized as a heatmap in R, with negative means set to zero for display. Exploratory outlier scans identified windows at or above the 99.9th percentile of F_ST_ within each regional comparison; windows exceeding this threshold in more than one comparison were termed recurrent outliers.

### (e) Genetic diversity and runs of homozygosity

For analyses requiring both variant and invariant sites, such as nucleotide diversity estimation in pixy [35], a separate all-sites dataset –comprising the same 50 curated individuals used for pca– was generated and filtered independently. Invariant records (ALT=’.’) were extracted by chromosome using GATK GenotypeGVCFs [28] with the --all-sites option prior to allele-frequency filtering, ensuring that variant and invariant datasets were derived from the same set of callable genomic positions. Only invariant records (ALT=’.’) were retained, with genotype depth <5 set to missing. SNPs were prepared from the genotype-filtered autosomal dataset before MAF filtering. After sample subsetting and recalculation of allele counts and missingness, biallelic SNPs were retained if missingness was ≤20% and 0 < AC < AN, without a minimum MAF threshold. Variant and invariant records were combined using bcftools, yielding 1,146,905,667 records: 1,145,122,611 invariant records and 1,783,056 SNPs. Genome-wide π and Tajima’s D were estimated in non-overlapping 50 kb windows across the 50 individuals treated as a single population; regional analyses used the same dataset. Genome-wide summaries were restricted to chromosome-level autosomal sequences (SUPER_1–SUPER_32). Both π and mean window-level Tajima’s D were summarized using callable-site weights, excluding windows without a defined estimate for the corresponding statistic.

To assess whether diversity and differentiation differed around centromeres, pixy windows were classified as overlapping annotated putative centromeric regions from Baalsrud et al. [23], falling within 1 Mb pericentromeric flanks, or belonging to the remaining autosomal background. For diversity summaries, windows with at least 1,000 callable sites were retained, and π and window-level Tajima’s D were averaged using callable-site weights. For pairwise F_ST_ the mean across regional comparisons was calculated per window and negative window means were set to zero before plotting and comparing the regional distributions with a Kruskal–Wallis test. F_ST_ windows were additionally grouped by distance from the nearest annotated centromere. Windows above the 99.9th-percentile F_ST_ threshold in at least one regional comparison were intersected with gene and repeat annotations. F_ST_ windows were additionally grouped by distance from the nearest annotated centromere.

Runs of homozygosity (ROHs) were identified from the MAF-filtered autosomal SNP dataset in 66 individuals using PLINK v1.9 [30] (Table S2). ROHs were detected using a sliding-window approach with a window size of 50 SNPs, allowing up to three heterozygous sites and ten missing genotypes per window. For a SNP to be assigned to a homozygous segment, at least 5% of overlapping windows were required to satisfy the homozygosity criteria. ROHs were defined as segments containing at least 50 consecutive SNPs, spanning a minimum length of 300 kb, with a minimum SNP density of one SNP per 50 kb and a maximum gap of 1 Mb between adjacent SNPs. Up to three heterozygous sites were permitted within an ROH to account for genotyping errors and sequencing artefacts. For each individual, genomic inbreeding (FROH) was estimated as the total length of all ROHs divided by the total length of the autosomal genome [36]. ROHs were additionally classified into short (0.3–1 Mb), intermediate (1–5 Mb), and long (>5 Mb) segments to distinguish signatures of older demographic processes from recent inbreeding.

### (f) Demographic history

Recent changes in effective population size (*N*_e_) were inferred using GONE [37–39], which estimates historical *N*_e_ from patterns of linkage disequilibrium. Analyses were conducted using a subset of autosomal microchromosomes selected based on chromosome homology and morphology. Because no recombination map is available for snowy owl, chromosome-specific recombination rates from chicken (*Gallus gallus*) were used as proxies [40,41]. To maximize biological comparability, only snowy owl microchromosomes exhibiting one-to-one synteny with chicken chromosomes and sharing comparable chromosome morphology were retained [23] (electronic supplementary materials, Text S6). The resulting chromosome set has a mean recombination rate of 6.63 cM/Mb.

GONE analyses were performed using genotype-filtered SNPs from the selected autosomal microchromosomes. No LD pruning was applied, because GONE infers effective population size from linkage disequilibrium among markers. The primary analysis used 24 high-depth individuals and polymorphic sites, and sensitivity analyses were performed using both the 24-individuals high-depth dataset and the full 64-individuals dataset with additional MAF thresholds of 0.005, 0.01, and 0.02 (electronic supplementary materials, Text S6 and Table S11). Individuals identified as contaminants and close relatives were excluded. Effective population size estimates were converted from generations to years applying a generation time of eight years [42].

## 3. Results

### (a) No evidence of population genetic structure

We sequenced the genomes of 68 snowy owls to a mean depth of 27x (ranging from 12x to 54x) (electronic supplementary materials, Table S2). Variant calling produced approximately 30 million SNPs. After the filtering steps (see Materials and Methods) we retained 1,324,912 SNPs in the MAF-filtered dataset. The autosomal PCA, based on the curated PCA-filtered sample set and LD-pruned SNPs, showed individuals from all sampling localities overlapping broadly, with no clear clustering by geographic region (electronic supplementary materials, Figure S2). The leading first four axes explained similar and modest proportions of genetic variance (2.09 - 2.15 %), without a dominant axis of geographic differentiation. To test whether this pattern depended on diploid genotype calls or filtering of the GATK dataset, we repeated the PCA using pseudohaploid genotypes generated independently with ANGSD, which is less sensitive to genotype-calling uncertainty in samples with variable sequencing depth. This second PCA did not reveal any clear separation among sampling regions either (electronic supplementary materials, Text S2 and Figure S2), yet the first axes explained only 2.20% and 2.19% of the variance, respectively.

The sex-linked PCAs did not provide robust evidence for sex-linked geographic structure (electronic supplementary materials, Text S2 and Figure S2). The Z-chromosome PCA showed a strong sex-associated axis, consistent with differences in sex chromosome ploidy and/or callability. The W-chromosome PCA was restricted to females; although Wrangel Island females showed partial separation, this pattern was confounded with sequencing batch, as Wrangel females sequenced in batch 4 formed the separated cluster whereas Wrangel females sequenced in batch 2 did not. Pairwise kinship analyses showed most individuals formed a single homogeneous cluster irrespective of sampling locality, further supporting the absence of strong population subdivision (Figure S3). A small cluster of three Fennoscandia individuals (FNM11, FNM12 and FNM15) had elevated genetic similarity relative to the remainder of the dataset (KING kinship coefficients = 0.125–0.153; electronic supplementary materials, Table S3). These individuals were likely close relatives, suggesting that this cluster reflected familial relatedness rather than population-level differentiation.

Genome-wide pairwise F_ST_ estimates among sampling localities were low, with SNP-weighted means ranging from −0.001237 to 0.00015 (Figure 1B; Table S5). Negative estimates were interpreted as no evidence of differentiation. Callable-site-weighted mean dXY ranged from 2.48 × 10^−4^ to 2.53 × 10^−4^, close to genome-wide nucleotide diversity, consistent with low absolute divergence among sampling regions (Table S5). The highest differentiation was observed between Greenland and New York (F_ST_ = 0.000155). These values indicate negligible genome-wide differentiation among sampled regions. Taken together, PCA, kinship analyses and pairwise F_ST_ estimates provide no evidence for substantial population structure across the sampled snowy owl range.

### (b) Genomic diversity in the snowy owl panmictic population

Genome-wide nucleotide diversity, – i.e., the callable-site-weighted mean pi – was 2.47 × 10^−4^ across approximately 1.10 Gb of callable autosomal sequence (Figure 2A). Mean proportion of autosomal heterozygous SNP genotypes was similar across individuals (mean = 0.175, MAF-filtered dataset), with only modest variation among sampling regions (electronic supplementary materials, Table S2). This SNP-based proportion is conditional on the filtered variant set and is not a per-base estimate of genome-wide diversity.

**Figure 2.**
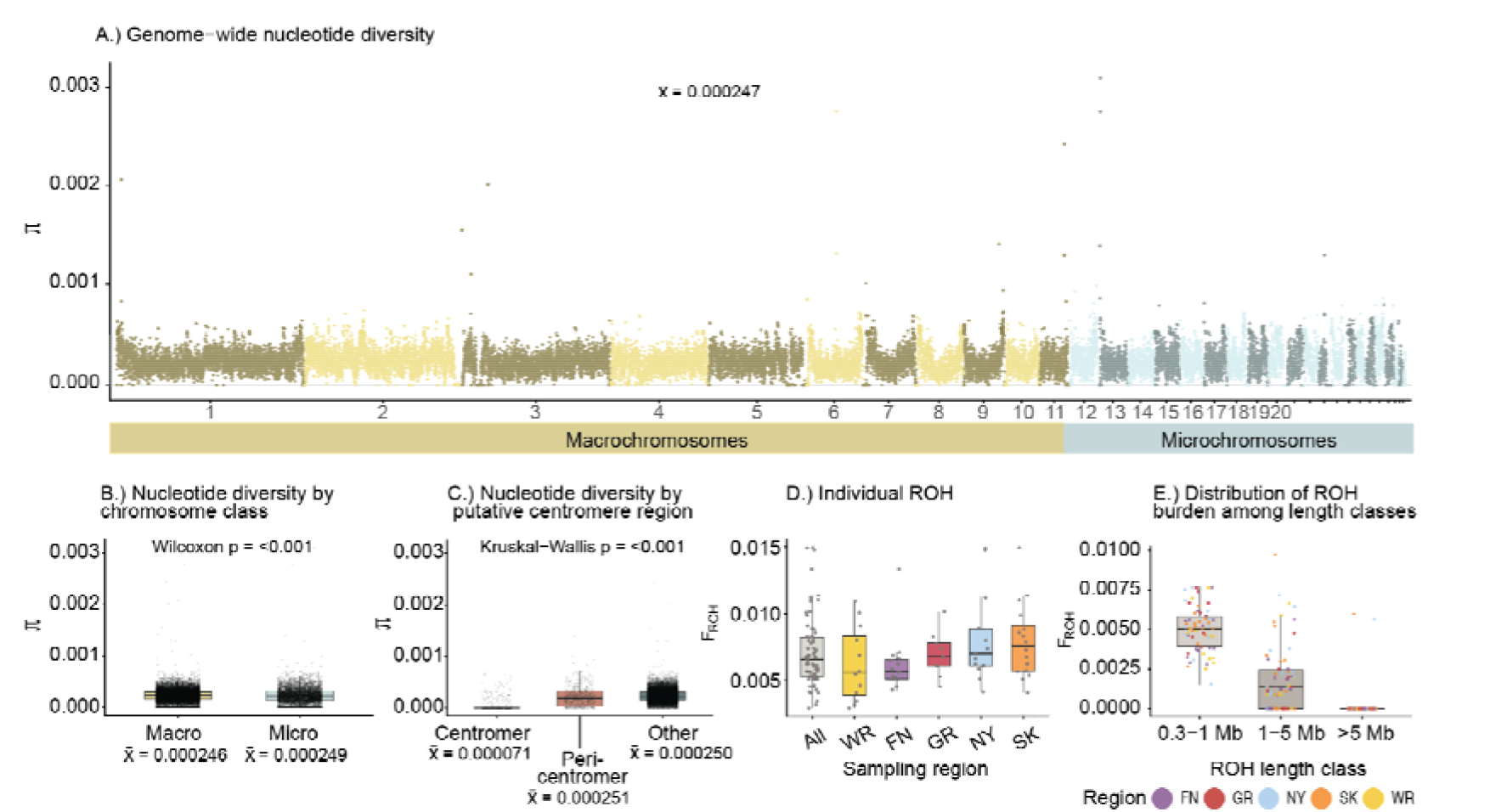
Genome-wide diversity and runs of homozygosity in snowy owls. A) Average genome-wide nucleotide diversity (π) calculated in non-overlapping 50 kb windows across autosomes in 50 curated individuals. Chromosomes are ordered by genomic position and colored by chromosome class. B, Distribution of π on macrochromosomes and microchromosomes. C, π in annotated putative centromeric regions, 1 Mb pericentromeric flanks, and the remaining genomic background. D, Individual genomic inbreeding coefficients estimated as FROH in 66 individuals, summarized by sampling region and across all individuals. E, Distribution of ROH burden among ROH length classes. Points represent individuals.

Nucleotide diversity varied across the genome, but was similar between microchromosomes and macrochromosomes (Figure 2B). Tajima’s D was negative (callable-site-weighted mean = −0.699), with slightly more negative values on microchromosomes than macrochromosomes (−0.721 and −0.692, respectively; electronic supplementary materials, Figurse S4 and S5). The negative mean indicates a relative excess of rare variants, but does not by itself identify the demographic or selective processes responsible.

Snowy owl chromosomes contain large satellite-rich centromeric regions, which have been hypothesized to evolve rapidly through processes such as centromeric drive [23]. We therefore compared diversity and differentiation in annotated centromeric regions, 1 Mb pericentromeric flanks, and the remaining genomic background. Nucleotide diversity was markedly reduced in centromeric regions compared with the rest of the genome, with weighted mean pi of 7.08 × 10^−5^, compared with 2.51 × 10^−4^ in pericentromeric windows and 2.50 × 10^−4^ in the remaining autosomal background (Figure 2C; electronic supplementary materials, Table S7). The weighted mean window-level Tajima’s D was slightly more negative in centromeric and pericentromeric windows (−0.749 and −0.813) than in the background (−0.696; electronic supplementary materials, Figure S7). Mean F_ST_ across regional comparisons showed no statistically detectable difference among the three categories after negative window means were set to zero (Kruskal–Wallis test, P = 0.969; electronic supplementary materials, Figure S8).

To investigate recent and historical inbreeding, we identified runs of homozygosity (ROH) across autosomal chromosomes. Across 66 individuals, we identified 946 ROH segments. Mean FROH across all individuals was low (0.0071), corresponding to an average of 8.34 Mb of the autosomal genome in ROH. Mean FROH was broadly similar among sampling regions, ranging from 0.0061 in Fennoscandia to 0.0082 in New York (Figure 2D). Most ROH segments were short: 90.0% were 0.3-1 Mb, 9.8% were 1-5 Mb, and only 0.2% were longer than 5 Mb (Fig 2E). The longest ROH we identified was 7.0 Mb (individual SKW03, electronic supplementary materials, Table S8). These results suggest historically small population size rather than inbreeding, although some individuals carried moderate ROH burdens (FROH=10-15%).

### (c) Recent trends in effective population size track climate changes

We used GONE [37–39] to infer recent changes in effective population size over the last ~200 generations, corresponding to 1,600 years for the snowy owl, which has a generation time of 8 years [42].In the primary GONE analysis, inferred Ne was highest early in the plotted trajectory, declining from approximately 50,000 individuals around AD 400 to a minimum of approximately 8,000-9,000 individuals around AD 1200 (Figure 3). Ne subsequently increased toward the present, reaching approximately 17,000-18,000 individuals in the most recent generations. This pattern is consistent with a long-term reduction in effective population size followed by partial recent recovery. Over the last few centuries, Ne fluctuated modestly around ~15,000-19,000 individuals, with no evidence for a sharp recent decline in the GONE trajectory, although point estimates should be interpreted cautiously because GONE reflects smoothed LD-based inference rather than census abundance.

**Figure 3.**
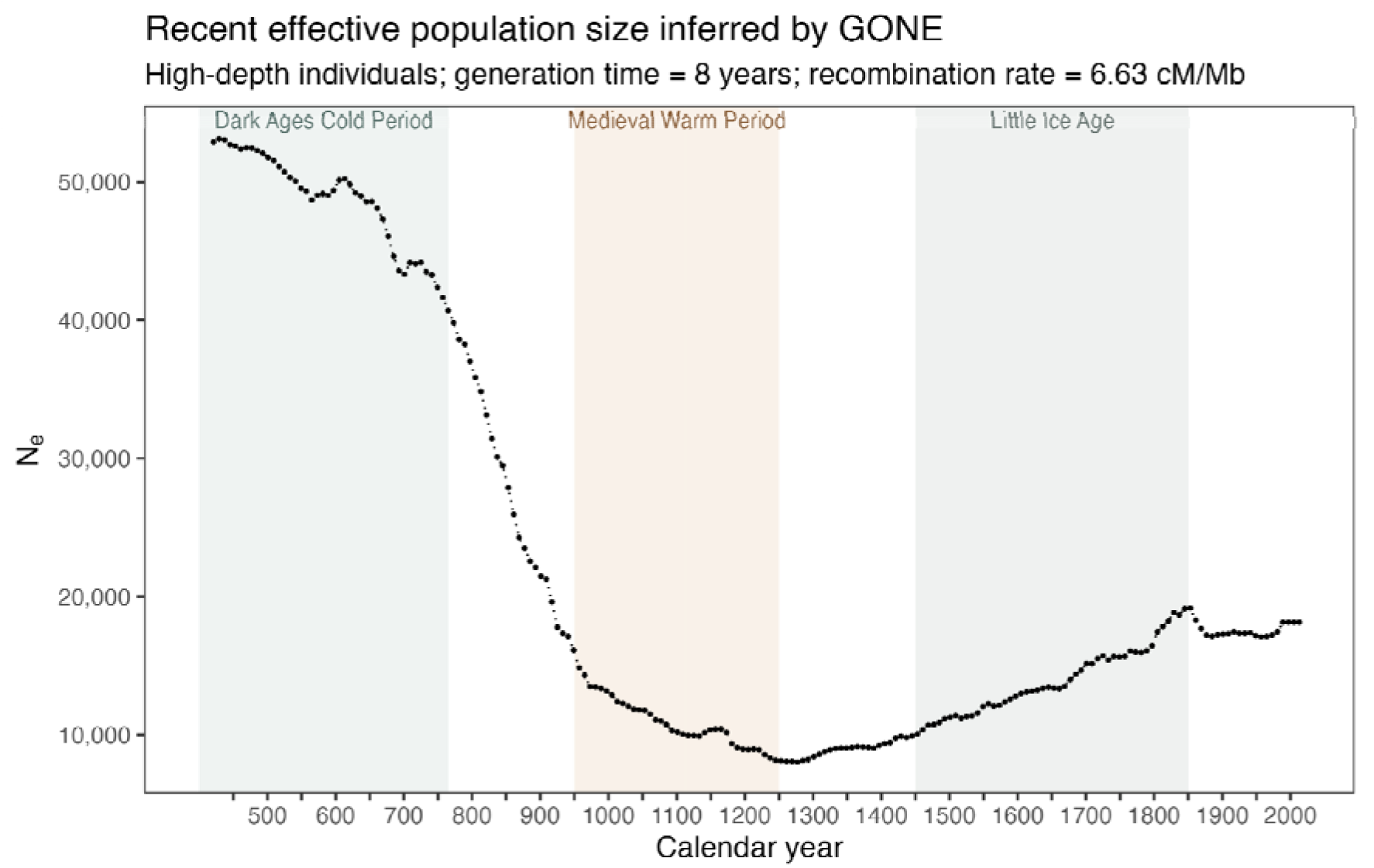
Recent demographic history of snowy owl. Effective population size (Ne) was inferred using GONE [37] from linkage disequilibrium using 24 high-depth individuals and a subset of syntenic autosomal microchromosomes (electronic supplementary materials, Table S10). Calendar years were calculated assuming a generation time of eight years. Shaded regions indicate the Dark Ages Cold Period (AD 400-765), the Medieval Warm Period (950-1250) and Little Ice Age (1450 to 1850) [56]. GONE results are shown for the most recent ~200 generations, where LD-based inference is expected to be most informative. The complete time series is provided in Online Data Table 7.

Sensitivity analyses indicated that GONE estimates were strongly affected by sample depth and rare-variant filtering. Analyses using all 64 individuals without sufficient rare-variant filtering produced implausibly high *N*_e_ estimates exceeding one million individuals, whereas the 24-individual high-depth dataset produced stable trajectories across filtering thresholds. Full-dataset analyses with MAF >= 0.01 or MAF >= 0.02 were more similar to the high-depth results, suggesting that the inflated estimates were driven by low-depth individuals and/or very rare variants. We therefore used the high-depth dataset as the primary demographic inference but retained the full sensitivity analysis in the electronic supplementary materials, Text S6.

## 4. Discussion

### (a) High connectivity and lack of population structure

Using whole-genome resequencing and a chromosome-level reference genome, this study provides the most comprehensive assessment of snowy owl population structure and genetic variation to date. Our results confirm and expand upon previous work based on mitochondrial markers and reduced-representation sequencing [9,10], demonstrating that snowy owls form a largely panmictic population across their circumpolar range. PCA, F_ST_ and relatedness analyses revealed no clustering by geographic origin, with the few apparent clusters attributable to either relatedness among sampled individuals or technical variation introduced during library preparation. The absence of detectable population structure is consistent with the ecology of snowy owls. As highly mobile predators that track spatiotemporal fluctuations in prey [3,5], snowy owls show low site fidelity to breeding grounds [43] and moderate to high fidelity to wintering grounds [13] while they disperse widely across continents. Although some individuals are known to return to the same general wintering areas in some years [11,13], this behavior does not appear sufficient to generate measurable genetic differentiation at a global scale, especially when considering the extent of annual breeding dispersal.

### (b) Genetic diversity and inbreeding

Snowy owls exhibit low genome-wide nucleotide diversity (2.47 × 10^−4^), which is low relative to most birds in other genomic datasets and consistent with patterns of reduced standing genetic diversity reported in other long-lived apex predators with modest global census sizes [44]. Our runs-of-homozygosity (ROH) results show limited evidence of recent inbreeding: ROH segments are short (<2.6 Mb), indicating primarily ancient or low-intensity background relatedness rather than ongoing mating among close relatives (Figure 2). These findings suggest that snowy owls currently are not inbred, yet have low standing genetic variation often associated with small or isolated populations [45–47]. Although many Arctic vertebrates exhibit signatures of inbreeding—such as Svalbard reindeer or the Arctic fox, where homozygous loss-of-function variants can directly reduce fitness [48,49]—snowy owls do not show equivalent genomic signs of inbreeding (Figure 2). This also contrasts with several threatened or lekking bird systems, including prairie-chickens [50], sage-grouse [51], black grouse [52] and kākāpō [53], in which fragmentation, bottlenecks or skewed reproductive success have been associated with elevated inbreeding, reduced diversity, or fitness-related genetic load. In snowy owls, high dispersal capacity and extensive gene flow across a largely panmictic range may help reduce the probability of repeated mating among close relatives, even when local abundance fluctuates.

### (c) Demographic history

Polar and Arctic species are expected to be especially sensitive to climatic warming because their distributions, prey dynamics, and breeding conditions are closely tied to snow, sea ice, and tundra ecosystems [54]. In line with this our reconstruction of effective population size (*N*_e_) over the past 1,600 years reveals a general decline, punctuated by climate-associated fluctuations (Figure 3). The trajectory begins during the Dark Ages Cold Period, a cooler and climatically disturbed interval commonly dated to approximately 400–765 CE in the Northern Hemisphere [55]. The most pronounced decrease coincided with the Medieval Warm Period (ca. 950–1250 CE), a period of regionally variable but often warmer conditions across parts of the North Atlantic and Arctic [56], suggesting that snowy owl populations may have been sensitive to warming conditions in northern ecosystems. Conversely, population size increased during the cooler conditions of the Little Ice Age (ca. 1450–1850 CE). These results align with long-term demographic reconstructions that place snowy owl populations on a declining trajectory since the Last Glacial Maximum [10]. Climate-associated demographic responses have also been inferred in other Arctic and polar taxa, including polar bears, Arctic foxes, collared lemmings, and Arctic char, supporting the broader view that cold-adapted species can show genomic demographic responses to past environmental change [57]. Our contemporary estimates place *N*_e_ at approximately 18,000, a figure that corresponds closely with global population estimates of 14,000–28,000 breeding adults from the International snowy owl Working Group [2]. Although *N*e and census size are not directly equivalent, this similarity suggests that snowy owls may maintain a comparatively high *N*e/*N* ratio [58]. This may reflect broad gene flow across a largely panmictic range, overlapping generations, and a breeding system with low variance in family size, all of which can help buffer effective population size against local demographic fluctuations

Our LD-based reconstruction (GONE) and the SFS-based trajectory (Stairway Plot 2) [10] show broadly concordant trends over the period of temporal overlap, with both methods showing an overall decline in *N*_e_. GONE inferred a sharper decrease around the Medieval Climate Anomaly and a subsequent partial recovery toward the Little Ice Age, whereas these recent fluctuations were not resolved by Stairway Plot 2. This difference is expected because LD-based approaches generally have greater resolution for recent demographic change, while SFS-based methods are better suited to deeper demographic history. Thus, the two analyses should be viewed as complementary

Our analyses were based on a single panmictic population, reducing the risk that cryptic structure biased the trajectory, [37], but snowy owls have overlapping generations, which could lead to underestimation of contemporary *N*_e_ [39]. Recombination rate also strongly influences *N*_e_ inference but is unknown in most non-model species and is therefore rarely accounted for [59]. Here, we inferred recombination rates for each snowy owl chromosome based on synteny with their chicken homologs [40,41], rather than relying on GONE’s default of 1 cM/Mb. As chicken recombination rates tend to be higher than those measured directly in other owls [41,60], this approach likely yields conservative rather than inflated *N*_e_ estimates. Therefore, the absolute population sizes may be underestimated, but the overall declining trajectory is unlikely to be an artefact of the recombination-rate assumption. Future work should refine this estimate using pedigree-based data.

The negative mean Tajima’s D indicates an excess of rare variants (electronic supplementary materials, Figurse S4 and S5). Such a pattern can arise after population expansion, during some bottleneck-and-recovery histories, or through selection [61]. It is therefore compatible with complex demographic history but does not independently confirm the decline inferred by GONE. The reduction in π in putative centromeric windows may reflect local processes such as reduced recombination and linked selection, including centromere drive, which would leave similar footprints in the genome as a strong selective sweep. Recurrent centromere drive could contribute to reduced diversity across multiple chromosomes, particularly around centromeres where recombination is suppressed. However, the genome-wide excess of rare variants and low nucleotide diversity cannot be attributed to centromere drive from these summary statistics alone. Moreover, as centromere annotations cover only a subset of chromosomes and these regions have confounding issues such as repetitive sequences, mapping uncertainty and differences in callability. Future studies testing whether centromeric variants show transmission bias during female meiosis could help evaluate the centromere-drive hypothesis in snowy owls.

### (d) Ecological and conservation implications

The ecological dependence of snowy owls on tundra-associated small mammal populations, such as lemmings and voles, provides a mechanistic explanation for the climate sensitivity observed in the demographic time series. Long-lived species with extended generation times are known to face disproportionately higher extinction risk under environmental change, since population-level impacts of reproductive failure compound across fewer, slower generational cycles [42]. Lemming and vole dynamics are tightly linked to snowpack conditions and seasonal thermal regimes, and disruptions to the high-amplitude fluctuations in their population sizes can substantially reduce reproductive output in snowy owls [8,62,63]. Thus, even a genetically healthy, well-connected species such as the snowy owl can experience demographic stress when prey availability becomes increasingly unpredictable [64–66]. In long-lived species with extended generation times it may take a long time before the effects of such stressors are reflected in population sizes, yet regional declines in snowy owls have already been reported. For example, in the latter half of the 19th century, the snowy owl commonly bred almost every 4 years in Norway. After 1909, breeding years appear to have become more irregular and generally fewer breedings [67–69]. Also, snowy owls once bred far south into alpine regions such as Hardangervidda until 1974 [69,70], but the species is now listed as critically endangered in Norway and Finland [71] and extinct in Sweden [72]. Similar downward trajectories are documented across other parts of the range [73,74]. Our results indicate that these declines are unlikely to be driven by inbreeding or population fragmentation; instead, they appear rooted in climatic warming and its cascading effects on prey and habitat. This distinction is important for conservation. Despite low genome-wide diversity, snowy owls show little evidence of recent inbreeding and a comparatively high recent effective population size, suggesting they do not face acute genetic viability issues, but they remain vulnerable to ecological and climatic pressures. Protecting this species therefore requires actions focused on habitat, prey dynamics, and climate mitigation, rather than genetic rescue or management of isolated demes. Indeed, the primary threat to snowy owl persistence is ecological: Arctic warming is altering lemming population cycles [75,76], destabilizing the prey base on which snowy owl reproduction depends [66], reshaping the tundra habitat required for nesting [77], and facilitating the northward expansion of boreal predators and competitors, further restructuring the tundra community [78,79].

## 5. Conclusion

By combining whole-genome resequencing, demographic inference, and assessments of heterozygosity, we show that snowy owls form a highly connected, and largely panmictic population with low genetic diversity. The species has experienced climate-linked demographic declines across both historical and contemporary time scales. Our results underscore that for snowy owls, conservation concern arises not from inbreeding, but low genetic diversity, compounded by continued population decline in a rapidly warming Arctic.

## Ethics

This study did not require ethical approval from a human subject or animal welfare committee. The data used in this study was obtained from museum collections, surveys and collection of shed feathers. No live animal subjects were involved in this study.

## Data accessibility

Sequencing data and metadata is available at ENA (PRJEB77068). Vcfs are available at Zenodo (reserved 10.5281/zenodo.22825317). Scripts and detailed description of data analysis is available at this Github repository: https://github.com/hellebaa/Snowy_Owl_population_genomics.

## Declaration of AI use

ChatGPT 5 and Codex have been used for text polishing of some paragraphs, as well as scripts. All content represents original work from the authors.

## Authors’ contributions

The study was conceived and designed by HTB, SB, and RN. Samples were provided and/or collected by KOJ, RS, IJØ, and TA in Fennoscandia; IM on Wrangel Island, Russia; JFT in the USA; and KLW in Canada. The Greenland samples originated from deceased birds recovered in Greenland by unknown finders and were obtained via Aarhus University; RS prepared the skins in Denmark and facilitated access to the material. Laboratory work, including DNA extraction, library preparation, and sequencing coordination, was carried out by HTB, ELGE, DAG, OSK, and SB. ELGE generated the initial analytical framework, performed early analyses, and prepared an initial manuscript draft. HTB and ELGE carried out the main population genomic analyses, with additional analytical input from DAG, SNKH, SB, and RN. HTB regenerated and curated the final analyses and figures. ELGE, DAG, and HTB wrote the manuscript, with DAG and HTB substantially revising and developing the final version. All authors contributed to interpretation of the results, reviewed the manuscript, and approved the final version.

## Acknowledgments

This work would not have been possible without the global network of researchers, conservationists, and scientists in ISOWG (International snowy owl Working Group). We thank Jan-Eric Hägerroth, Henrik Lind, and Tom McDonald for collecting feathers in Sweden and the US. We also want to thank Tore Slagsvold for early brainstorming of this project and Lydia Smith for her advice and assistance during library preparation. The computations were performed on resources provided by Sigma2 - the National Infrastructure for High-Performance Computing and Data Storage in Norway. Sequencing was performed at the NSC (The Norwegian Sequencing Centre) and the OMRF (Oklahoma Medical Research Foundation) Clinical Genomics Center.

## Supplementary Materials and Methods

Contains the following supplementary sections:

### Supplementary Text S1. Sampling and variant datasets

#### Sampling and sequencing

We obtained samples from five locations across the snowy owl range (Figure 1): Saskatchewan, Canada (n = 17), Wrangel Island, Russia (n = 12), Fennoscandia (n = 16), Greenland (n = 10), and New York, USA (n = 13). Samples were collected between 1996 and 2020 as part of ongoing and previous research and monitoring efforts by project collaborators. The dataset included both adult and juvenile individuals and comprised material from live birds, including feathers and blood feathers, as well as muscle tissue from deceased birds. Complete sample metadata are provided in Table S1.

**Supplementary Table S1.** Sample metadata and sequencing summary. Metadata for all snowy owl individuals included in the study, including sample identifiers, sampling region, country/locality, collection year, observed sex (sex_field), and age. Abbreviations: M = male, F = female, NA = unavailable or unknown; MU = muscle, BL = blood, BF = blood feather, FE = feather; T = included/true, F = excluded/false.

| sample_id | sample_short_id | individual_id | country | location | pop_id | Collection year | Sex field | age | tissue |
| --- | --- | --- | --- | --- | --- | --- | --- | --- | --- |
| FNM01_9-X42_S17_L002 | FNM01 | FNM01 | NA | Fennoscandia | FN | 2011 | NA | Nestling | BF |
| FNM02_10-X43_S20_L002 | FNM02 | TB00878 | NA | Fennoscandia | FN | 2011 | NA | Nestling | BF |
| FNM03_11-X44_S23_L002 | FNM03 | TB00405 | NA | Fennoscandia | FN | 2011 | NA | Nestling | BF |
| FNM04_12-X45_S26_L002 | FNM04 | TB00407 | NA | Fennoscandia | FN | 2011 | NA | Nestling | BF |
| FNM05_13-X46_S29_L002 | FNM05 | TB00402 | NA | Fennoscandia | FN | 2011 | NA | Nestling | BF |
| FNM06_14-X47_S32_L002 | FNM06 | FNM06 | NA | Fennoscandia | FN | 2011 | NA | Nestling | BF |
| FNM07_15-X48_S35_L002 | FNM07 | TB00410 | NA | Fennoscandia | FN | 2011 | NA | Nestling | BF |
| FNM08_merged_X1 | FNM08 | BA40507 | NA | Fennoscandia | FN | 2015 | NA | Nestling | BL |
| FNM09_merged_X2 | FNM09 | FNM09 | NA | Fennoscandia | FN | 2015 | NA | Nestling | BL |
| FNM10_merged_X3 | FNM10 | FNM10 | NA | Fennoscandia | FN | 2015 | NA | Nestling | BL |
| FNM11_merged_X4 | FNM11 | FNM11 | NA | Fennoscandia | FN | 2015 | NA | Nestling | BL |
| FNM12_5-X5_S119_L004 | FNM12 | FNM12 | NA | Fennoscandia | FN | 2015 | NA | Nestling | BL |
| FNM13_merged_X6 | FNM13 | FNM13 | NA | Fennoscandia | FN | 2015 | NA | Nestling | BL |
| FNM15_merged_X8 | FNM15 | BA40510 | NA | Fennoscandia | FN | 2015 | NA | Nestling | BL |
| FNM16_16-X49_S37_L002 | FNM16 | BA40505 | NA | Fennoscandia | FN | 2015 | NA | Nestling | BL |
| FNM17_17-X50_S18_L002 | FNM17 | TB00412 | NA | Fennoscandia | FN | 2011 | NA | Nestling | BF |
| GRL01_merged_X11 | GRL01 | 2438 | Greenland | Eastern Greenland | GR | 1996 | F | Adult | MU |
| GRL02_merged_X12 | GRL02 | 2445 | Greenland | Eastern Greenland | GR | 2008 | F | Juvenile | MU |
| GRL03_merged_X13 | GRL03 | 2439 | Greenland | Eastern Greenland | GR | 2000 | F | Juvenile | MU |
| GRL04_merged_X14 | GRL04 | 2440 | Greenland | Eastern Greenland | GR | 2001 | M | Juvenile | MU |
| GRL05_merged_X15 | GRL05 | 2437 | Greenland | Eastern Greenland | GR | 2006 | F | Adult | MU |
| GRL06_merged_X16 | GRL06 | 2444 | Greenland | Eastern Greenland | GR | 2001 | M | Juvenile | MU |
| GRL07_merged_X17 | GRL07 | 2442 | Greenland | Eastern Greenland | GR | 1998 | M | Juvenile | MU |
| GRL08_merged_X18 | GRL08 | 2446 | Greenland | Eastern Greenland | GR | 1998 | M | Juvenile | MU |
| GRL09_merged_X19 | GRL09 | 2441 | Greenland | Eastern Greenland | GR | 2000 | M | Juvenile | MU |
| GRL10_merged_X20 | GRL10 | 2443 | Greenland | Eastern Greenland | GR | 2001 | F | Juvenile | MU |
| NY01_DAG-BS-1_S1 | NY01 | Unbanded (dead) | USA | New York State | NY | 2021 | M | Second-year | FE |
| NY02_DAG-BS-2_S2 | NY02 | 0699-03116 | USA | New York State | NY | 2020 | F | After hatch-year | FE |
| NY03_DAG-BS-3_S3 | NY03 | 0829-00946 | USA | New York State | NY | 2019 | F | Second-year | FE |
| NY04_DAG-BS-4_S4 | NY04 | 0829-00947 | USA | New York State | NY | 2020 | F | After first-year | FE |
| NY05_DAG-BS-5_S5 | NY05 | 0829-00948 | USA | New York State | NY | 2020 | F | After third-year | FE |
| NY06_DAG-BS-6_S6 | NY06 | 0289-00949 | USA | New York State | NY | 2020 | F | After first-year | FE |
| NY07_DAG-BS-7_S7 | NY07 | 0829-00950 | USA | New York State | NY | 2020 | F | Third-year | FE |
| NY08_DAG-BS-8_S8 | NY08 | 0289-00951 | USA | New York State | NY | 2020 | F | After third-year | FE |
| NY09_DAG-BS-9_S9 | NY09 | 0829-00952 | USA | New York State | NY | 2020 | F | After hatch-year | FE |
| NY10_DAG-BS-10_S10 | NY10 | 0829-00953 | USA | New York State | NY | 2020 | F | After third-year | FE |
| NY11_DAG-BS-11_S11 | NY11 | 1238-04099 | USA | New York State | NY | 2021 | M | Second-year | FE |
| NY12_DAG-BS-12_S12 | NY12 | NY200822 | USA | New York State | NY | 2020 | F | Hatch-year | FE |
| NY13_DAG-BS-13_S13 | NY13 | NY200818 | USA | New York State | NY | 2020 | M | After second-year | FE |
| SKW01_merged_X21 | SKW01 | 2415 | Canada | Saskatchewan | SK | 2010 | F | Adult | MU |
| SKW02_merged_X22 | SKW02 | 2416 | Canada | Saskatchewan | SK | 2005 | F | Juvenile | MU |
| SKW03_merged_X23 | SKW03 | 2417 | Canada | Saskatchewan | SK | 2013 | F | Adult | MU |
| SKW04_merged_X24 | SKW04 | 2419 | Canada | Saskatchewan | SK | 2009 | M | Sub adult | MU |
| SKW05_merged_X25 | SKW05 | 2420 | Canada | Saskatchewan | SK | 2010 | M | Sub adult | MU |
| SKW06_merged_X26 | SKW06 | 2421 | Canada | Saskatchewan | SK | 2008 | F | Adult | MU |
| SKW07_merged_X27 | SKW07 | 2422 | Canada | Saskatchewan | SK | 2004 | F | Adult | MU |
| SKW08_merged_X28 | SKW08 | 2423 | Canada | Saskatchewan | SK | 2008 | M | Juvenile | MU |
| SKW09_merged_X29 | SKW09 | 2424 | Canada | Saskatchewan | SK | 2013 | M | Juvenile | MU |
| SKW10_merged_X30 | SKW10 | 2425 | Canada | Saskatchewan | SK | 2006 | F | Juvenile | MU |
| SKW11_merged_X31 | SKW11 | 2426 | Canada | Saskatchewan | SK | 2008 | F | Juvenile | MU |
| SKW12_merged_X32 | SKW12 | 2427 | Canada | Saskatchewan | SK | 2008 | F | Juvenile | MU |
| SKW13_merged_X33 | SKW13 | 2429 | Canada | Saskatchewan | SK | 2006 | F | Juvenile | MU |
| SKW14_merged_X34 | SKW14 | 2430 | Canada | Saskatchewan | SK | 2007 | F | Juvenile | MU |
| SKW15_merged_X35 | SKW15 | 2431 | Canada | Saskatchewan | SK | 2005 | F | Juvenile | MU |
| SKW16_merged_X36 | SKW16 | 2432 | Canada | Saskatchewan | SK | 2008 | F | Juvenile | MU |
| SKW17_merged_X37 | SKW17 | 2436 | Canada | Saskatchewan | SK | 2002 | M | Juvenile | MU |
| WRG01_1-WRG1_S28_L001 | WRG01 | WRG01 | Russia | Wrangel Island | WR | 2014 | NA | Adult | FE |
| WRG02_2-WRG2_S33_L001 | WRG02 | WRG02 | Russia | Wrangel Island | WR | 2014 | NA | Adult | FE |
| WRG03_3-WRG3_S39_L001 | WRG03 | WRG03 | Russia | Wrangel Island | WR | 2014 | NA | Adult | FE |
| WRG05_4-WRG5_S4_L001 | WRG05 | WRG05 | Russia | Wrangel Island | WR | 2015 | NA | Adult | FE |
| WRG06_21-X54_S30_L002 | WRG06 | WRG06 | Russia | Wrangel Island | WR | 2014 | NA | Adult | FE |
| WRG08_5-WRG8_S10_L001 | WRG08 | WRG08 | Russia | Wrangel Island | WR | 2016 | NA | Adult | FE |
| WRG09_6-WRG9_S15_L001 | WRG09 | WRG09 | Russia | Wrangel Island | WR | 2017 | NA | Adult | FE |
| WRG10_7-WRG10_S19_L001 | WRG10 | WRG10 | Russia | Wrangel Island | WR | 2018 | NA | Adult | FE |
| WRG11_8-WRG11_S24_L001 | WRG11 | WRG11 | Russia | Wrangel Island | WR | 2019 | NA | Adult | FE |
| WRG32_18-X51_S21_L002 | WRG32 | WRG32 | Russia | Wrangel Island | WR | 2014 | NA | Adult | FE |
| WRG46_19-X52_S24_L002 | WRG46 | WRG46 | Russia | Wrangel Island | WR | 2014 | NA | Adult | FE |
| WRG48_20-X53_S27_L002 | WRG48 | WRG48 | Russia | Wrangel Island | WR | 2014 | NA | Adult | FE |

DNA from blood and tissue samples was extracted using Qiagen’s DNeay Blood and Tissue kit, according to protocol, except the amount of proteinase K was doubled to improve lysis. For the feather samples, approximately 1 cm of the calamus from up to 6 feathers was used for DNA extractions, using Qiagens DNeasy Blood & Tissue kit and following the protocol for nail/hair/feathers. The amount of 1 M DTT was increased from 20 µl to 50 µl and the amount of proteinase K was tripled. The Qubit dsDNA Quantitation, Broad Range kit (Thermo Fisher) was used to determine the concentrations of each sample. Genomic libraries were built using TruSeq DNA Nano preparation kit (Illumina) and sequencing was carried out using Illumina NovaSeq 6000 (NovaSeq S4). Eight feather samples from Russia had too low DNA concentrations to use the TruSeq kit, therefore, libraries were built with the ThruPLEX preparation kit. Library preparations and sequencing were carried out by the Norwegian Sequencing Centre for all samples except for the 13 samples from New York state, which were sequenced at Oklahoma Medical Research Foundation.

#### Variant calling and filtering

Adapter sequences and low-quality bases were removed using Trimmomatic v0.39 [24], retaining reads with a minimum length of 50 bp. Filtered reads were aligned to the chromosome-level snowy owl reference genome [23] using BWA-MEM v0.7.17 [25]. Alignments were sorted and indexed using Samtools v1.17 [80], and PCR duplicates were identified and removed using Picard v3.0.0 [81]. Alignment statistics were generated with Qualimap v2.3 [26] and summarized with MultiQC v1.14 [27] (Table S2). Alignment QC indicated high mapping success (88-99%), with 61 samples having >99% aligned reads. Downstream filtering and sample exclusions were therefore based primarily on genotype-level QC metrics, including autosomal depth, missingness, heterozygosity, genomic sex concordance, contamination screening, and relatedness.

Variant calling followed GATK Best Practices (GATK v4.4; [28,82]). Variants were first called independently in each sample using HaplotypeCaller in GVCF mode. Individual GVCFs were subsequently combined and jointly genotyped using GenotypeGVCFs. SNPs and indels were separated using SelectVariants, and only SNPs were retained for downstream analyses.

Initial hard filtering was performed using GATK VariantFiltration. Variants were removed if they met any of the following criteria: QD < 2.0, FS > 60.0, SOR > 3.0, MQ < 40.0, MQRankSum < −12.5, ReadPosRankSum < −8.0, or BaseQRankSum < −6.0, when the corresponding annotations were present. Only PASS variants were retained.

Sample identifiers were standardized and variants were split into autosomal, Z chromosome, and W chromosome datasets. Because sex chromosomes require sex-specific treatment, downstream population genomic analyses were conducted using autosomes only. Genotype-level filtering was then applied using bcftools, with genotypes having depth <5× or genotype quality <20 converted to missing data. Following genotype filtering, allele counts (AC), allele numbers (AN), minor allele frequencies (MAF), and site missingness (F_MISSING) were recalculated using bcftools.

Autosomal SNPs were retained if they had a minor allele frequency ≥0.02 and missingness ≤20%. No upper depth filter was applied, as exploratory analyses indicated that previous depth-based filtering removed a substantial number of high-quality sites. The resulting dataset contained 1,324,912 autosomal SNPs and was used for for the MAF-filtered SNP analyses, including relatedness and ROH. A separate no-MAF dataset was generated for all pixy analyses (Text S3), and GONE datasets were prepared separately as described in Text S6.

#### Contamination, relatedness and depth filtering

During quality control, two individuals (GRL04 and SKW02) showed elevated heterozygosity and inconsistent sex chromosome signals, indicative of sample contamination or library mix-up. These individuals were excluded from downstream analyses. Additional sample exclusions based on missingness, other QC flags and relatedness were applied separately for different downstream analyses (Figure S1; Table S2).

Library-level inspection confirmed that contamination was present across sequencing libraries for these individuals, suggesting the issue originated prior to sequencing. These samples were therefore removed, and analyses were conducted on the remaining individuals.

To further assess data quality and identify potential sample contamination, we evaluated allele balance at heterozygous sites across all individuals. For each sample, we calculated the ratio of reference to total allele depth (AD) at heterozygous sites and summarized both the mean and standard deviation of this ratio. There was no correlation between depth and heterozygosity (Figure S1).

In a diploid genome, allele balance at heterozygous sites is expected to be centered around 0.5. The majority of samples showed mean allele balance values tightly centered around 0.5 (mean ≈ 0.50) with low variance (standard deviation ≈ 0.08–0.13), consistent with high-quality genotype calls.

Two individuals (GRL04 and SKW02) deviated markedly from this pattern, showing shifted mean allele balance (~0.56) and substantially elevated variance (~0.20). These patterns are characteristic of mixed DNA samples or contamination. In addition, these individuals exhibited elevated heterozygosity and inconsistent sex chromosome coverage patterns. Based on these combined lines of evidence, GRL04 and SKW02 were excluded from all downstream analyses. For pixy, further curation retained 50 individuals without high-missingness or contamination-review flags and excluded close relatives except FNM12, the retained representative of the related Fennoscandia group (Table S2).

The curated pixy set differed from the PCA, ROH and GONE datasets; the corresponding inclusion flags are reported in Table S2. A focused rare-variant quality check and sample-sensitivity analysis are described in Text S3.

**Figure S1.**
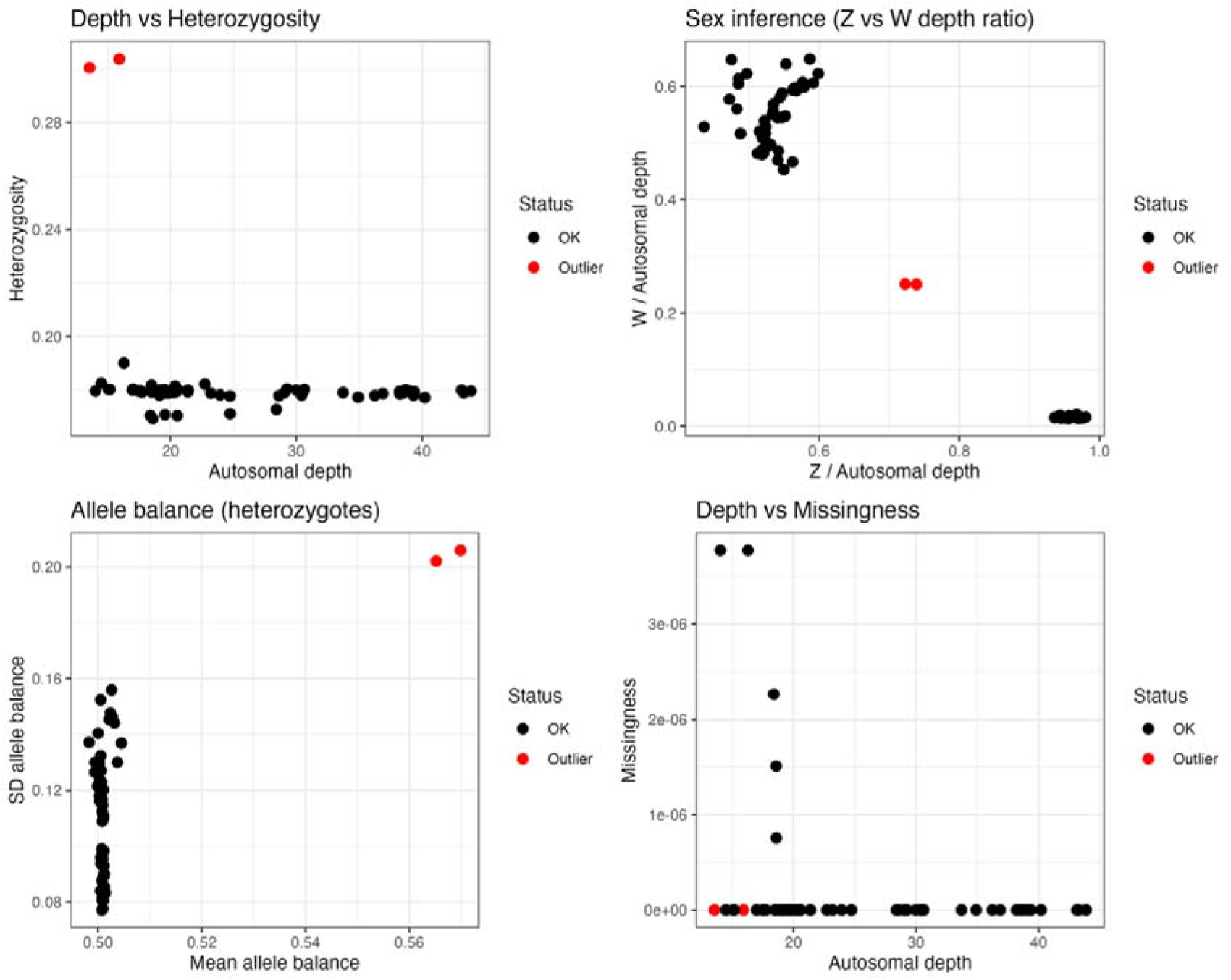
Quality-control metrics used to evaluate sequencing and variant quality prior to final filtering. (A) Mean sequencing depth versus observed heterozygosity for each individual. (B) Mean allele balance of heterozygous genotypes plotted against the standard deviation of allele balance for each individual. Individuals with balanced heterozygous allele ratios cluster around an allele balance of 0.5 with low variance, whereas deviations may indicate sequencing or genotyping artefacts. (C) Ratio of mean sequencing depth on the Z and W chromosomes, used to infer genetic sex and identify potential sample inconsistencies. (D) Mean sequencing depth versus genotype missingness for each individual. These quality-control metrics were used to identify contaminated or low-quality samples and to guide the selection of genotype- and site-filtering thresholds for downstream analyses.

**Supplementary Table S2.**
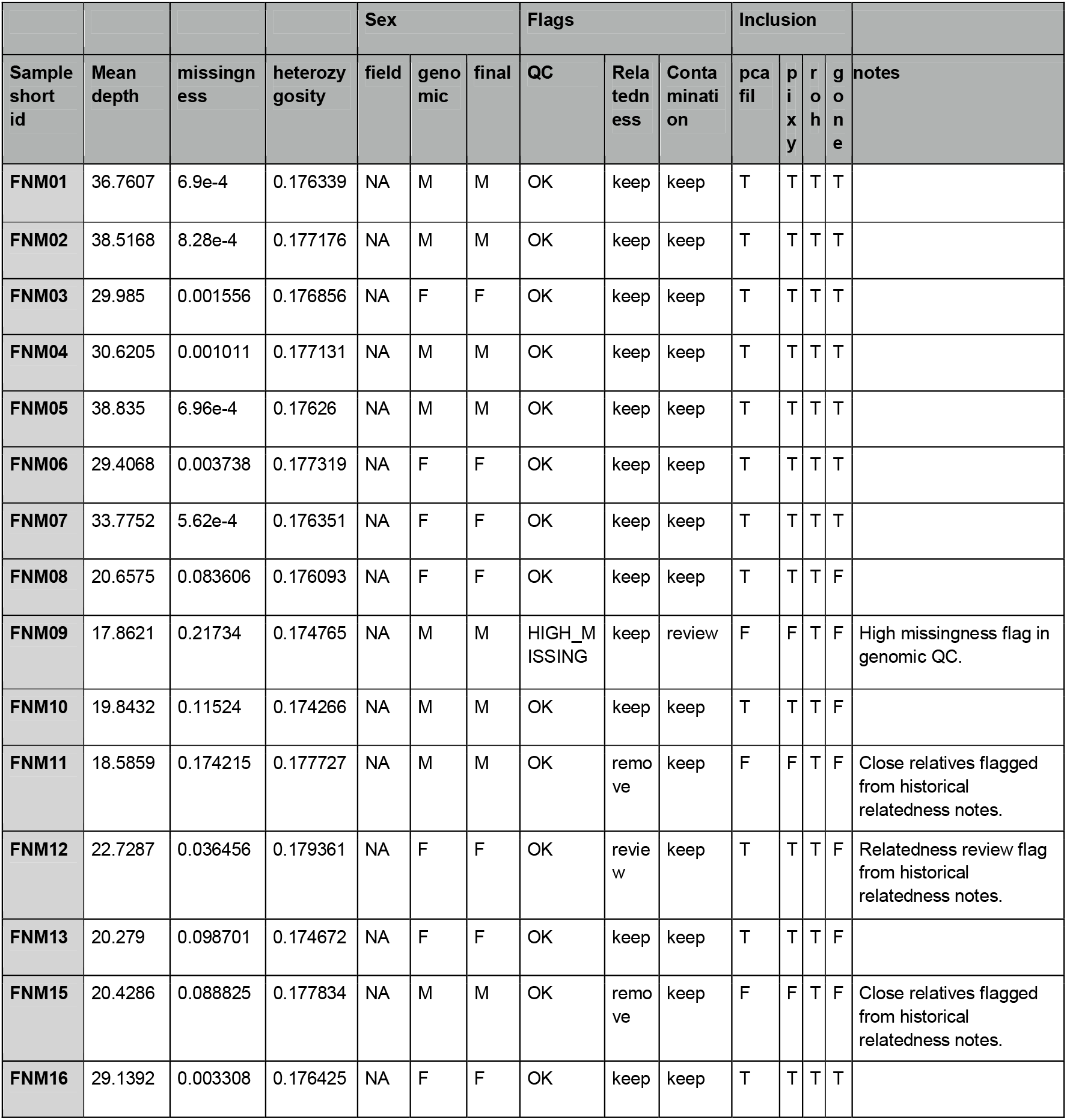
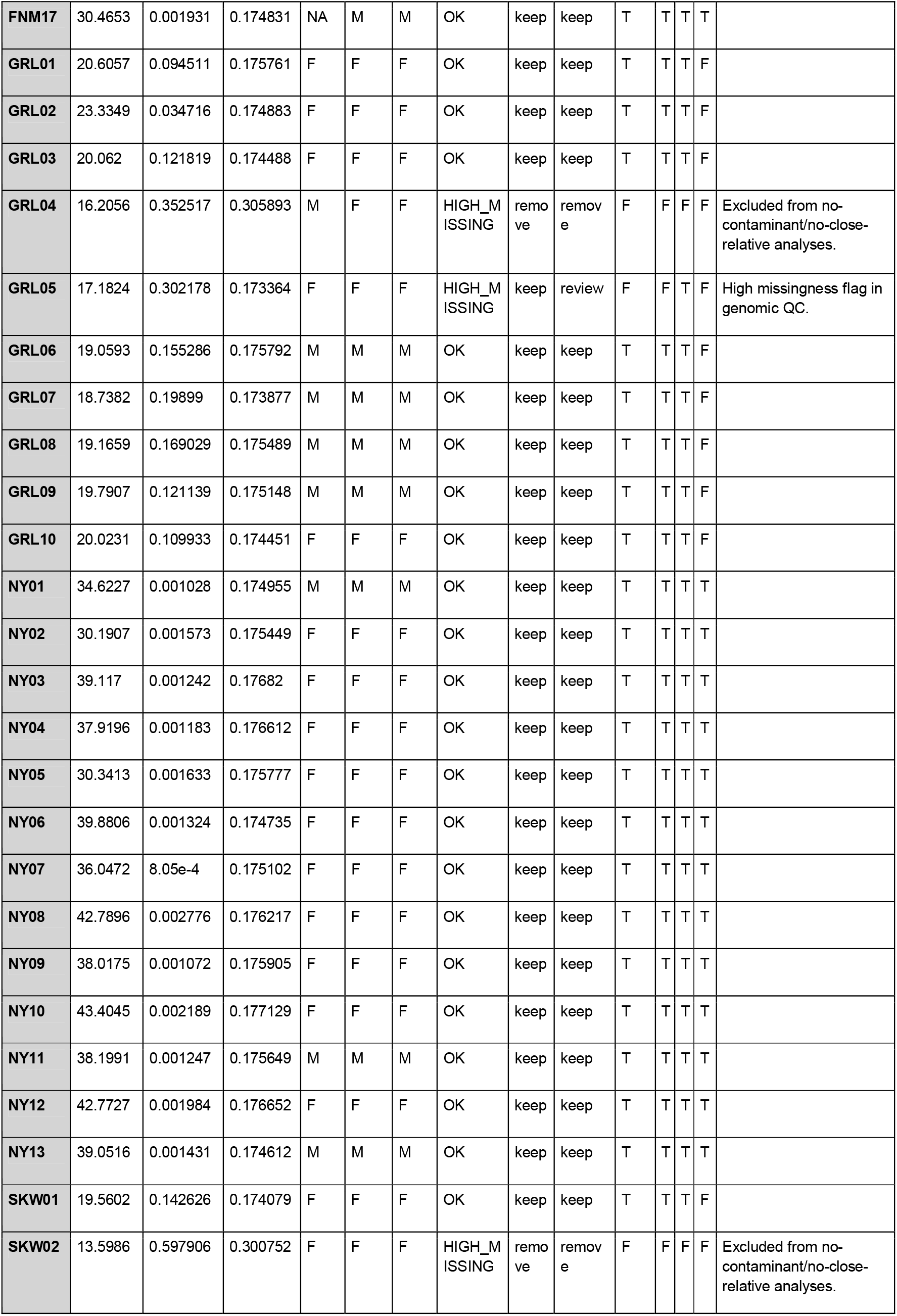
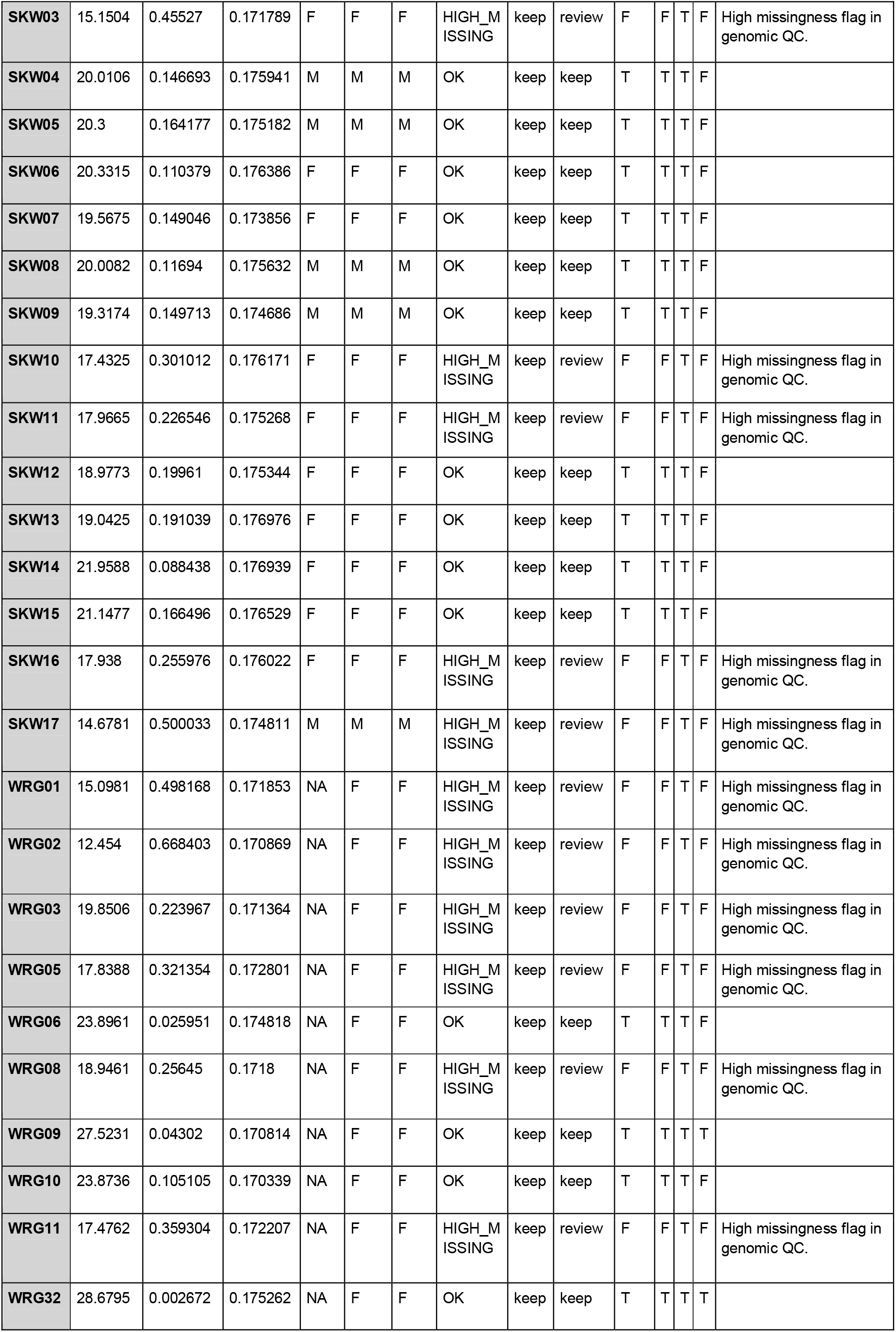
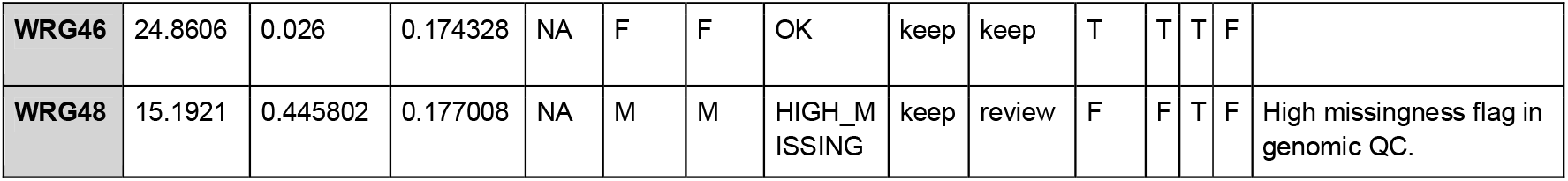
Sequencing and genotype QC summary. Sample-level sequencing and genotype QC metrics, including autosomal mean depth, missingness, heterozygosity (calculated as the proportion of non-missing autosomal SNP genotypes that were heterozygous for each individual), observed/field sex, genomic sex inferred from sex-chromosome depth, final interpreted sex, QC flags, relatedness and contamination decisions, and inclusion in downstream analyses. Abbreviations: M = male, F = female, NA = unavailable/unknown; T = included/true, F = excluded/false.

**Supplementary Table S3.**
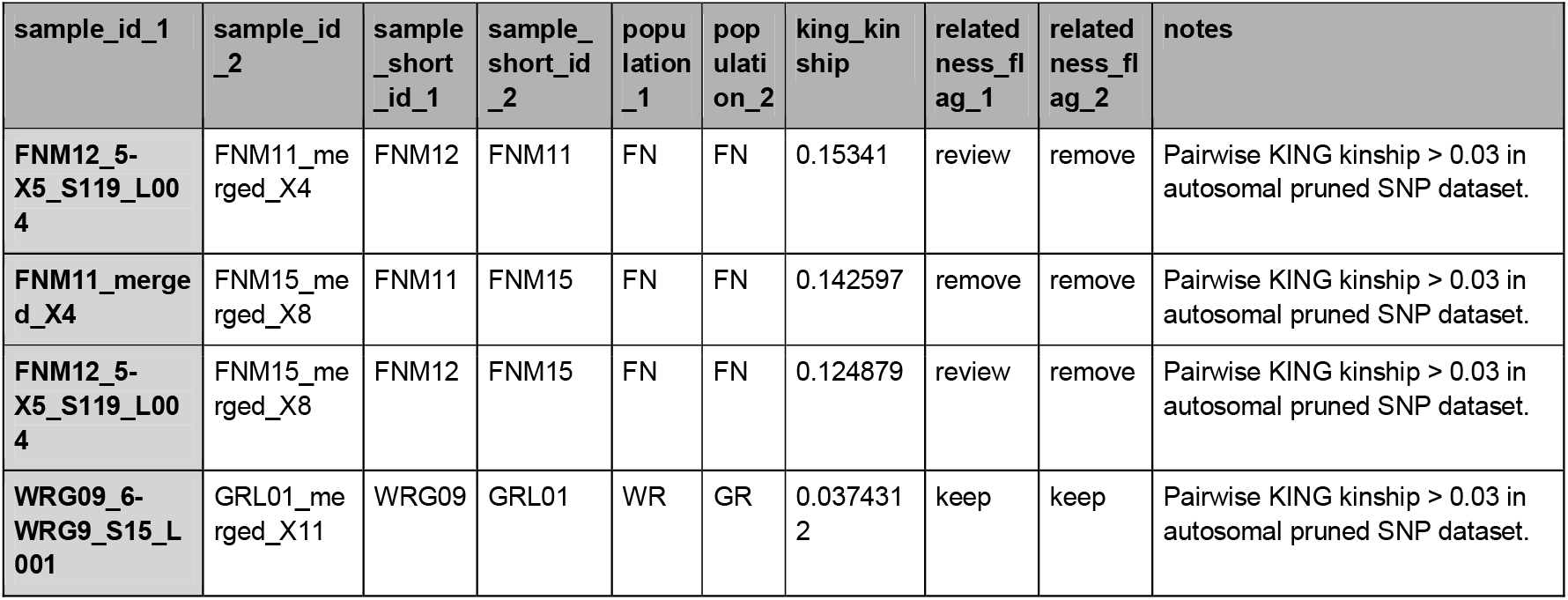
Pairwise KING kinship estimates above the reporting threshold. Pairwise KING kinship estimates greater than 0.03 calculated from the autosomal pruned SNP dataset. The full all-pair KING kinship table is provided as an online data table.

### Supplementary Text S2. Population structure

We used principal component analysis (PCA) and KING kinship estimates to assess broad population structure and to identify samples whose placement was likely influenced by contamination, sequencing quality, or close relatedness. Genotype-based PCA was performed in PLINK/PLINK2 [30,31] using the filtered autosomal GATK SNP dataset. Prior to PCA, variants were LD-pruned in PLINK using a 50-SNP window, 10-SNP step size, and r2 threshold of 0.2. KING kinship coefficients were estimated from the same LD-pruned autosomal genotype dataset using the PLINK2 implementation of KING-robust kinship estimation [31,83]. Because PCA filtering is described in the Methods, we summarize here only the sample-level decisions relevant to Figs. S2 and S3.

The autosomal PCA analyses shown in Fig. S2 used a curated 50-individuals sample set (also used for pixy). Five samples were excluded from the final visualization after inspection of the all-sample PCA, heterozygosity, missingness, contamination, and relatedness summaries: GRL04_merged_X14 and SKW02_merged_X22, which were independently flagged for contamination/high missingness; FNM11_merged_X4 and FNM15_merged_X8, which belonged to a closely related Fennoscandia group; and WRG01_1-WRG1_S28_L001, which behaved as an outlier in PCA and QC summaries. FNM12_5-X5_S119_L004 was retained as the representative individual from the related Fennoscandia group. The retained sample comprised 13 individuals from Fennoscandia, eight from Greenland, 13 from New York, 11 from Saskatchewan and five from Wrangel Island.

The final GATK autosomal PCA included 50 individuals and 564,409 SNPs, after MAC ≥3 filtering and LD pruning. As an independent comparison, we also performed PCA from ANGSD-derived pseudohaploid genotype data [33]. The ANGSD pseudohaploid autosomal PCA used the same curated PCA-filtered sample set and retained biallelic polymorphic sites with minor allele count >= 3, resulting in 50 individuals and 39,231 sites. This analysis provides a conservative comparison to the genotype-based GATK PCA because pseudohaploid genotype sampling and rare-variant filtering reduce axes driven by private alleles, missingness, or genotype uncertainty. In both autosomal PCAs, individuals showed extensive overlap among sampling regions, with no discrete geographic clusters.

We also generated ANGSD pseudohaploid PCAs for the Z and W chromosomes as diagnostic analyses. The Z-chromosome PCA included both sexes and showed a strong sex-associated axis, consistent with differences in sex chromosome ploidy and/or callability rather than geographic structure. The W-chromosome PCA was restricted to females.

Although Wrangel Island females showed partial separation in the W-linked PCA, this pattern was confounded with sequencing batch: Wrangel females sequenced in round 04 formed the separated cluster, whereas Wrangel females sequenced in round 02 did not. We therefore treat the sex-linked PCAs as QC diagnostics rather than evidence for sex-linked population structure.

The KING heatmap (Fig. S3) confirmed that most pairwise kinship values were low. The main exception was the Fennoscandia group FNM11, FNM12, and FNM15, which showed elevated relatedness relative to the rest of the dataset. These results support treating the sampled snowy owls as weakly structured for genome-wide diversity analyses, while excluding contaminated, outlier, or closely related individuals from analyses where those samples could disproportionately influence the results.

**Figure S2.**
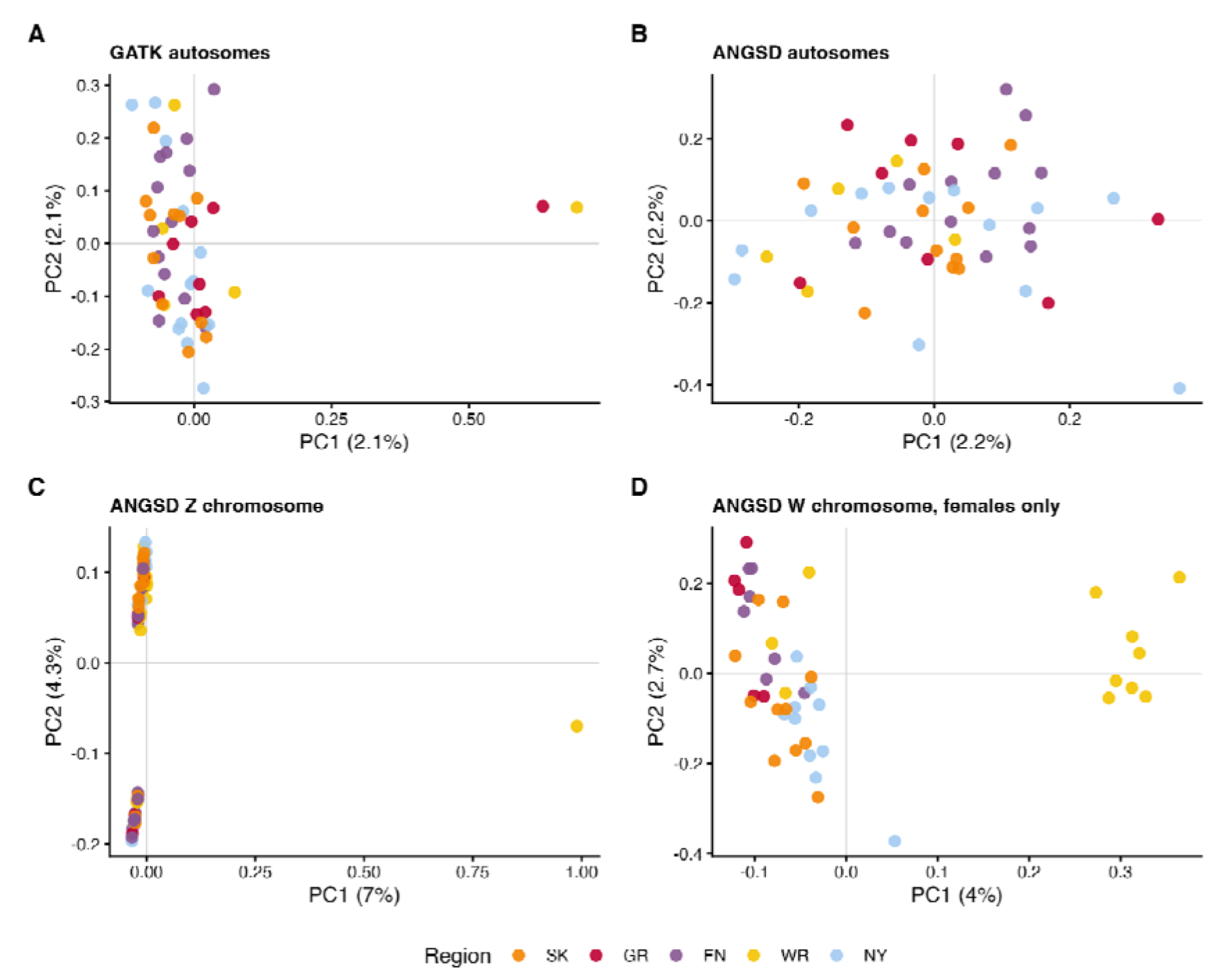
PCA of autosomal and sex-linked variation. Based on 50 curated individuals after removal of samples failing final QC, relatedness, or contamination filters (A) Genotype-based PCA from the filtered autosomal GATK SNP dataset with minor allele count ≥ 3 and LD pruning. (B) ANGSD pseudohaploid autosomal PCA using minor allele count ≥ 3. (C) ANGSD pseudohaploid Z chromosome PCA. (D) ANGSD pseudohaploid W chromosome PCA including female samples only. Points are coloured by sampling region.

**Supplementary Table S4.** PCA dataset summaries. Summary of PCA datasets used to assess population structure, including input data type, sample and site filters, number of retained samples, number of retained sites or SNPs where available, PC1/PC2 variance explained, and recommended figure use.

| dataset | input_type | sample_filter | site_filter | n_samples | n_sites_or_snps | pc1_variance_percent | pc2_variance_percent |
| --- | --- | --- | --- | --- | --- | --- | --- |
| **GATK autosomes, filtered** | diploid SNP genotypes | 50 individuals retained after QC and relatedness filtering; one representative retained per closely related group. | Existing Run A markers: MAC $\geq 3$ and LD pruning in 66 individuals | 50 | 564409 | 2.145 | 2.102 |
| **ANGSD pseudohaploid autosomes** | pseudohaploid alleles | 50 individuals retained after QC and relatedness filtering; one representative retained per closely related group. | Biallelic sites; MAC $\geq 3$ recalculated in 50 individuals | 50 | 39231 | 2.203 | 2.187 |
| **ANGSD pseudohaploid Z** | pseudohaploid alleles | 66 non-contaminated individuals | Biallelic sites; MAC $\geq 1$ | 66 | 3259198 | 6.97 | 4.26 |
| **ANGSD pseudohaploid W** | pseudohaploid alleles | 44 genetically female individuals | Biallelic sites; MAC $\geq 1$ | 44 | 29517 | 3.971 | 2.709 |

**Figure S3.**
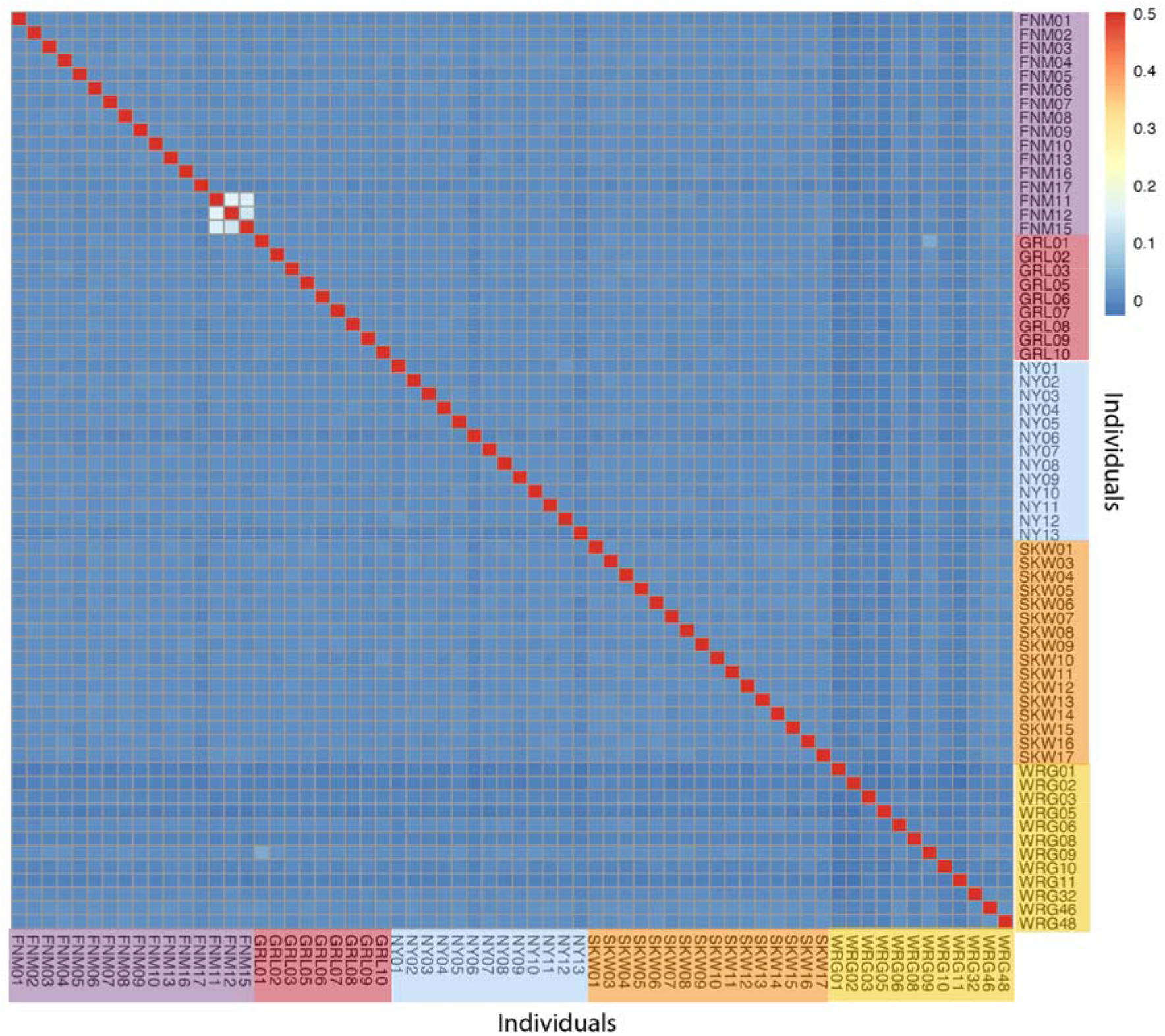
Heatmap of pairwise genetic similarity estimated using the KING kinship coefficient. Pairwise genetic similarity among individuals estimated using the KING kinship statistic calculated from linkage disequilibrium-pruned autosomal SNPs, colored according to legend. Individuals are ordered by sampling locality. Most individuals formed a single homogeneous cluster, consistent with limited population structure. A small cluster of three Fennoscandia individuals (FNM11, FNM12 and FNM15) showed elevated genetic similarity relative to the remainder of the dataset.

### Supplementary Text S3. Genome-wide diversity and differentiation

#### Generation of callable invariant sites for pixy analyses

To obtain unbiased estimates of nucleotide diversity, invariant callable sites were incorporated following the recommendations of the pixy workflow. Chromosome-level VCFs produced by GATK GenotypeGVCFs using the --all-sites option were used as the starting point. Invariant positions were identified as records lacking an alternate allele (ALT=’.’).

Sample identifiers were standardized and both variant and invariant VCFs were subset to the 50 curated individuals (curated50, Table S2). Invariant records were retained without site-level QUAL or RGQ filtering; individual invariant genotypes with DP <5 were set to missing. For the SNP component, the genotype-filtered autosomal dataset was used before the MAF ≥0.02 filter. After sample subsetting, AC, AN, MAF and F_MISSING were recalculated. Biallelic SNPs with F_MISSING ≤0.20 and 0 < AC < AN were retained, without a minimum MAF threshold.

Invariant records were concatenated across autosomes and combined with the curated no-MAF SNP dataset using bcftools. The resulting all-sites VCF contained 1,145,122,611 invariant records and 1,783,056 SNPs, yielding 1,146,905,667 records before pixy missing-data accounting. Summaries across chromosome-level autosomes comprised 23,489 non-overlapping 50 kb windows and 1,101,113,709 callable sites. π was defined in 23,485 windows and Tajima’s D in 22,256 windows; weights for each statistic included only windows with a defined estimate. All final pixy summaries, including F_ST_, dXY and outlier scans, used this curated50 no-MAF dataset.

To assess sensitivity to sample composition, we compared no-MAF analyses on chromosome 10 (SUPER_10) using 66 individuals, the same set without WRG01 (65 individuals), and the curated 50-individual set. Weighted mean π was 2.67 × 10⁻□, 2.38 × 10⁻□ and 2.36 × 10⁻□, respectively; corresponding weighted mean Tajima’s D was −1.846, −0.754 and −0.715. The large shift after removing WRG01 showed that this sample strongly influenced the rare-variant signal. In the curated set, singleton and common heterozygotes on chromosome 10 both had median alternate-allele balance of 0.50 and median genotype quality of 99; median depths were 27 and 26, respectively. These diagnostics supported use of the curated set but did not constitute a genome-wide test of variant accuracy.

#### Diversity measures

Tajima’s D was negative on average genome-wide (weighted mean = −0.699), with slightly higher values on macrochromosomes than microchromosomes (−0.692 vs. −0.721; Wilcoxon rank-sum test, P < 0.001) (Figure S5 and S6).

Pairwise differentiation among sampling regions was effectively zero (Table S5; Figure S6). Across all regional comparisons, mean weighted F_ST_ was −0.000246, consistent with weak population structure (Figure S7). Genome scans did not reveal strong, broad peaks suggestive of pronounced local adaptation. However, 187 unique windows above the pair-specific 99.9th-percentile threshold were identified in at least one comparison, of which 35 were recurrent outliers in more than one comparison. All 187 windows overlapped repeat annotations, particularly LINE/CR1 elements, and 122 overlapped annotated genes. These overlaps do not demonstrate enrichment without comparison to the genomic background. Many high-FST windows involved Wrangel Island comparisons, which warrant caution because only five individuals were retained from that region. The empirical outliers do not by themselves establish local adaptation.

**Supplementary Table S5.** Pairwise regional differentiation and divergence. Window-based pairwise differentiation and divergence among sampling regions, including the number of windows, SNP-weighted mean F_ST_, median F_ST_, upper F_ST_ quantiles, maximum F_ST_, site-weighted mean dXY, and median dXY. All estimates use the curated50 no-MAF dataset; n_windows refers to the F_ST_ windows passing the minimum of 10 SNPs. Negative FST estimates are retained here but set to zero in Figure 1B.

| comparison | populati<br>on_1 | populat<br>ion_2 | n_wind<br>ows | mean_fst_w<br>eighted_by_s<br>nps | median_fs<br>t | q95_fst | q99_fst | max_fst | mean_dxy_<br>weighted_b<br>y_sites | median_dxy |
| --- | --- | --- | --- | --- | --- | --- | --- | --- | --- | --- |
| GR_vs_NY | GR | NY | 22015 | 1.55e-4 | -0.002408 | 0.033883 | 0.058456 | 0.149261 | 2.5e-4 | 2.35e-4 |
| FN_vs_GR | FN | GR | 22015 | 4.3e-5 | -0.002431 | 0.033665 | 0.058963 | 0.166939 | 2.5e-4 | 2.35e-4 |
| FN_vs_SK | FN | SK | 22015 | -4.5e-5 | -0.002248 | 0.028513 | 0.04966 | 0.11985 | 2.52e-4 | 2.37e-4 |
| GR_vs_SK | GR | SK | 22015 | -1.07e-4 | -0.002881 | 0.036397 | 0.061656<br>9999999<br>99997 | 0.150607 | 2.53e-4 | 2.38e-4 |
| FN_vs_NY | FN | NY | 22015 | -1.45e-4 | -0.002121 | 0.025603 | 0.0452 | 0.120172 | 2.48e-4 | 2.34e-4 |
| NY_vs_WR | NY | WR | 22015 | -1.8e-4 | -0.003322 | 0.045964 | 0.079834 | 0.248503 | 2.5e-4 | 2.34e-4 |
| NY_vs_SK | NY | SK | 22015 | -2.33e-4 | -0.002386 | 0.028189 | 0.048863 | 0.115409 | 2.52e-4 | 2.37e-4 |
| FN_vs_WR | FN | WR | 22015 | -2.52e-4 | -0.003569 | 0.04669 | 0.080611 | 0.221327 | 2.51e-4 | 2.35e-4 |
| SK_vs_WR | SK | WR | 22015 | -4.61e-4 | -0.003882 | 0.048347 | 0.081624 | 0.369905 | 2.53e-4 | 2.38e-4 |
| GR_vs_WR | GR | WR | 22015 | -0.001237 | -0.005087 | 0.053907 | 0.091461 | 0.291899 | 2.51e-4 | 2.35e-4 |

**Figure S4.**
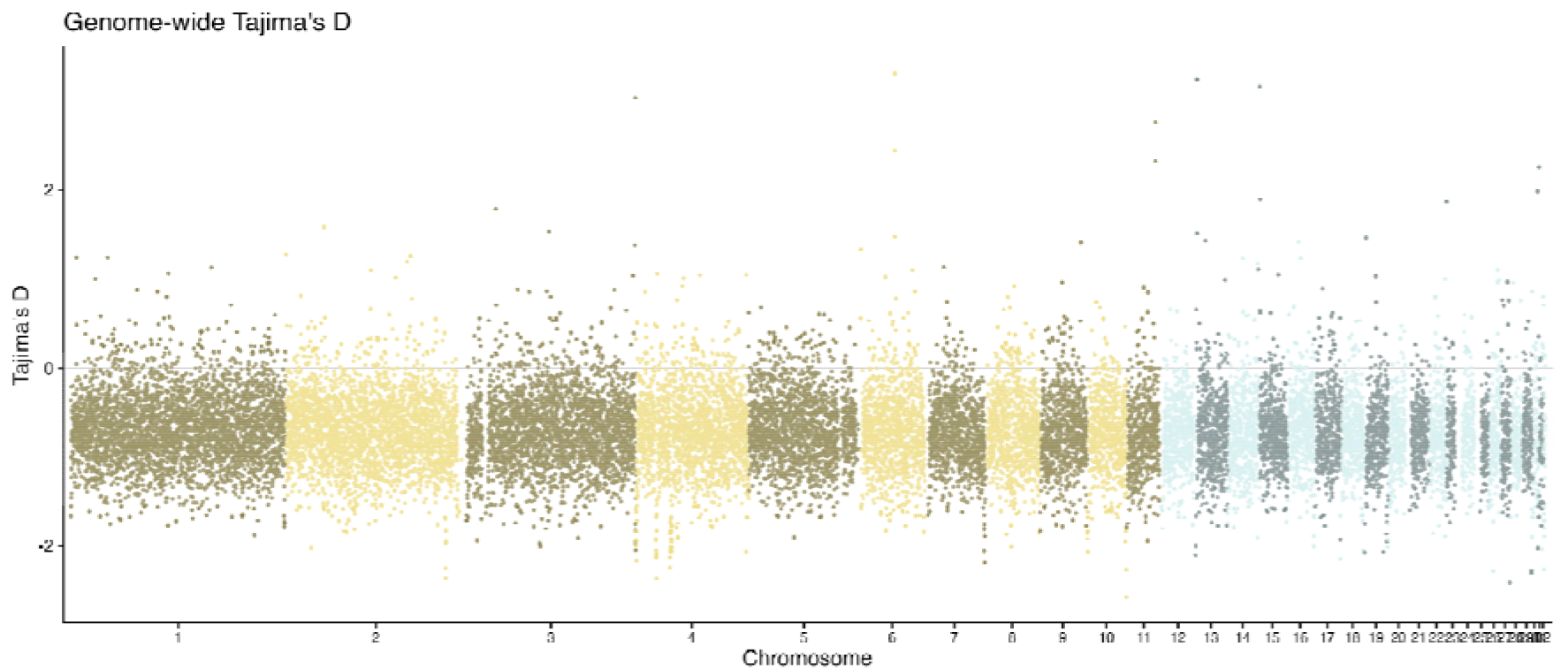
Genome-wide Tajima’s D calculated in non-overlapping 50 kb autosomal windows across the 50 curated individuals. Chromosomes are ordered by genomic position with macrochromosomes in yellow and microchromosomes in blue.

**Figure S5.**
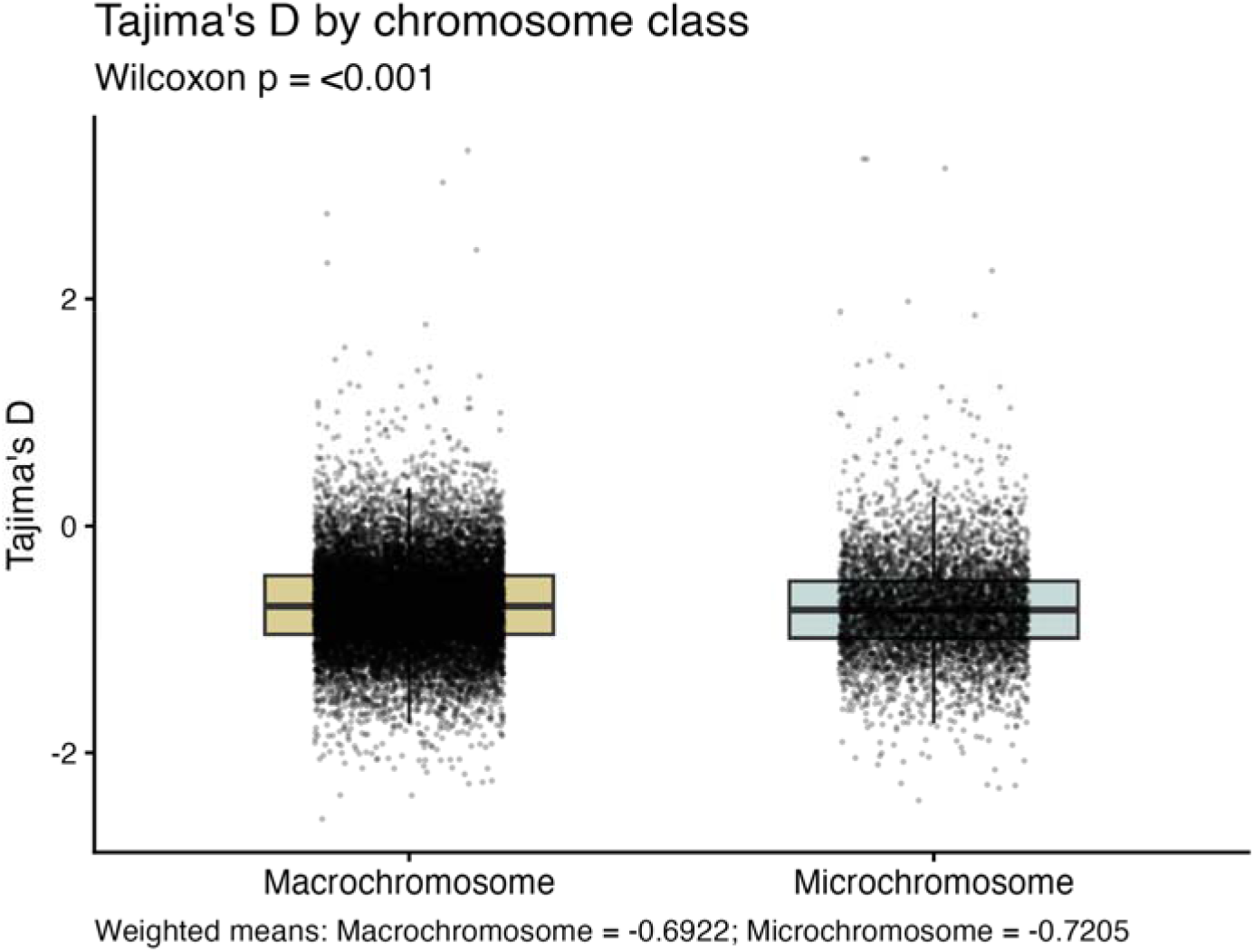
Window-level Tajima’s D compared between macrochromosomes and microchromosomes. Boxes show chromosome-window distributions; points show individual windows with defined estimates from the curated50 dataset. The Wilcoxon test compares chromosome-class distributions.

**Table S6.** Genome-wide diversity by chromosome class. Nucleotide diversity and Tajima’s D summarized separately for macrochromosomes and microchromosomes, including number of windows, callable sites, unweighted mean, callable-site-weighted mean, and median values for the curated50 dataset. Window counts and callable-site totals are statistic-specific and exclude windows without defined estimates.

| metric | chromosome_class | n_windows | callable_sites | mean_unweighted | mean_weighted_by_sites | median |
| --- | --- | --- | --- | --- | --- | --- |
| pi | Macrochromosome | 17381 | 822964037 | 2.4e-4 | 2.46e-4 | 2.38e-4 |
| pi | Microchromosome | 6104 | 278149672 | 2.35e-4 | 2.49e-4 | 2.26e-4 |
| Tajima_D | Macrochromosome | 16692 | 809034930 | -0.692849 | -0.692191 | -0.710674 |

**Figure S6.**
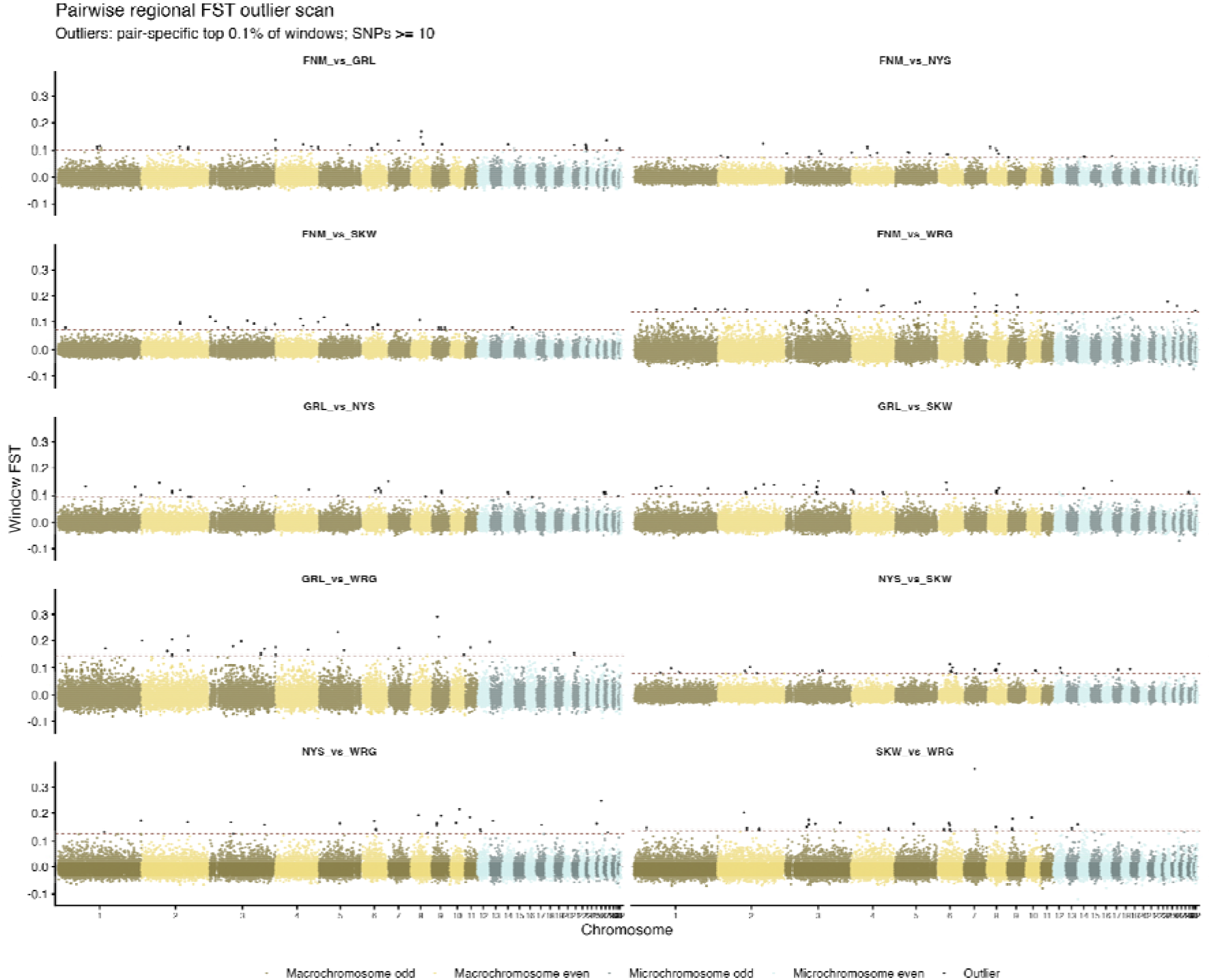
Genome-wide F_ST_ scans across non-overlapping 50 kb autosomal windows in the curated50 dataset. Windows contain at least 10 SNPs; highlighted outliers meet the pair-specific 99.9th-percentile threshold

### Supplementary Text S4. Centromeres and genomic features

Centromere positions are based on Baalsrud et al [23].

**Figure S7.**
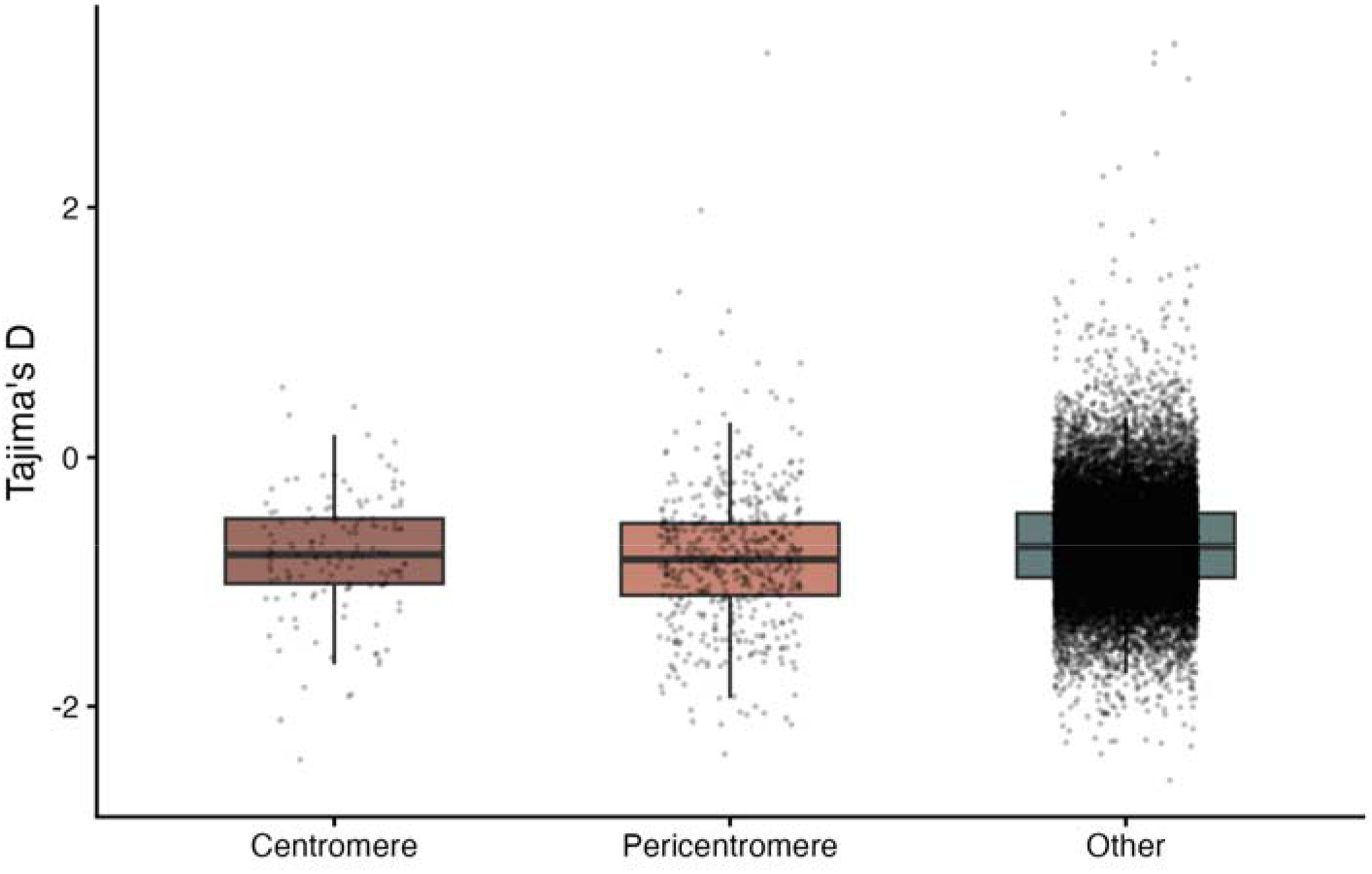
Tajima’s D in centromeric regions. Tajima’s D estimated in 50 kb autosomal windows and grouped by genomic region: annotated centromeric regions, 1 Mb pericentromeric flanks, and the remaining autosomal background. Centromeric regions showed lower Tajima’s D than the genomic background.

**Table S7.** Diversity in centromeric, pericentromeric, and background regions. Nucleotide diversity and Tajima’s D summarized for windows classified as putative centromere, pericentromere, or genomic background, including callable-site threshold, number of windows, callable sites, unweighted mean, callable-site-weighted mean, median, minimum, and maximum values for the curated50 dataset. n_windows_with_value gives the number of defined estimates; weighted means exclude undefined values. The callable_sites column sums sites across all retained windows and is not necessarily the denominator used for the Tajima’s D weighted mean.

| metric | centromere_region | min_no_sites | n_windows | n_windows_with_value | callable_sites | mean_unweighted | mean_weighted_by_sites | median | min | max |
| --- | --- | --- | --- | --- | --- | --- | --- | --- | --- | --- |
| avg_pi | Centromere | 1000 | 884 | 884 | 20444630 | 3.5e-5 | 7.1e-5 | 0 | 0 | 7.94e-4 |
| avg_pi | Pericentromere | 1000 | 653 | 653 | 27237949 | 2.28e-4 | 2.51e-4 | 1.99e-4 | 0 | 0.002714 |
| avg_pi | Other | 1000 | 21929 | 21929 | 1053420720 | 2.48e-4 | 2.5e-4 | 2.41e-4 | 0 | 0.003061 |
| tajima_d | Centromere | 1000 | 884 | 142 | 20444630 | -0.784364 | -0.749491 | -0.781662 | -2.421583 | 0.56265 |
| tajima_d | Pericentromere | 1000 | 653 | 529 | 27237949 | -0.807645 | -0.813357 | -0.820232 | -2.377165 | 3.240916 |
| tajima_d | Other | 1000 | 21929 | 21585 | 1053420720 | -0.696922 | -0.696281 | -0.715996 | -2.585398 | 3.309718 |

**Figure S8.**
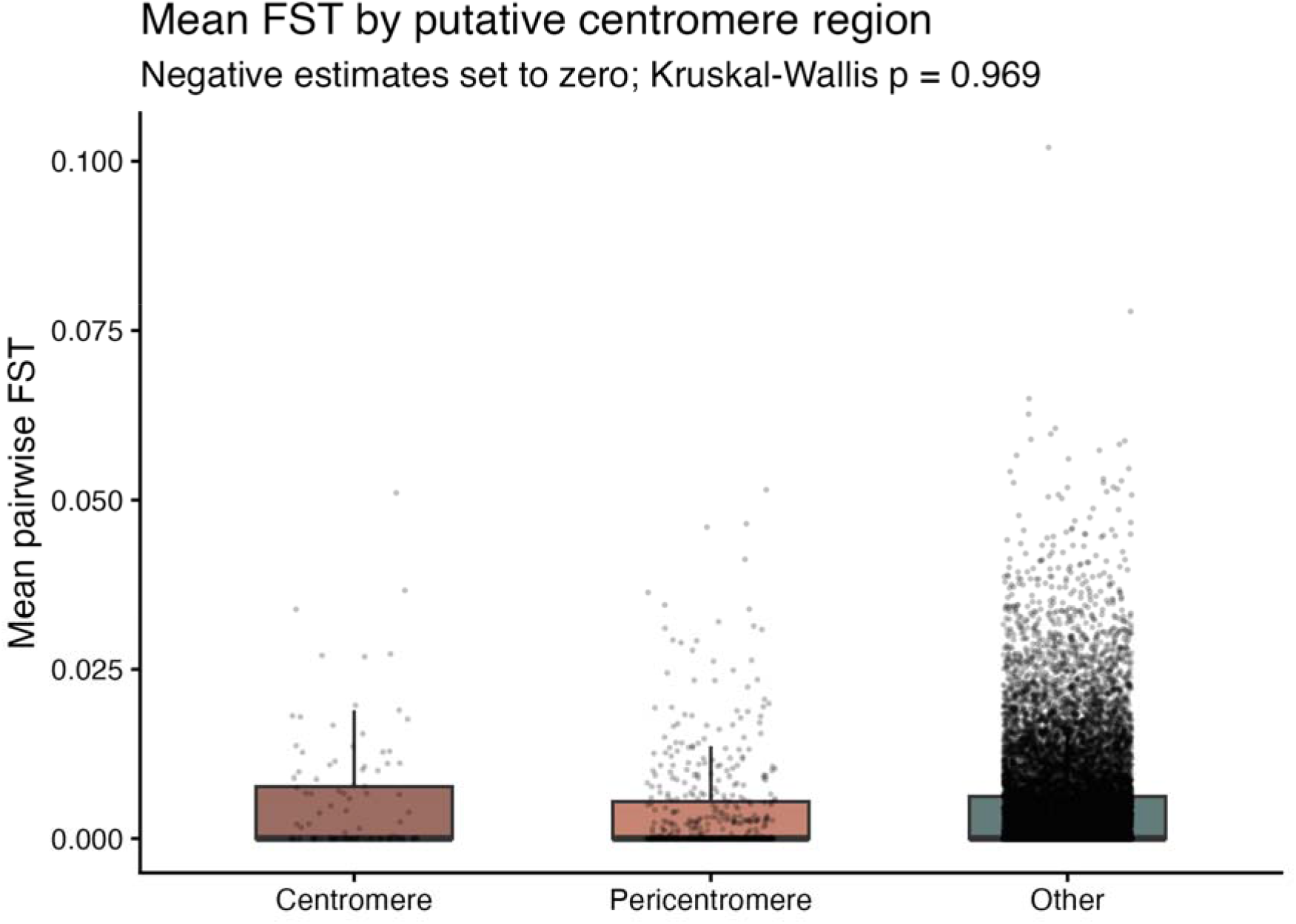
Genetic differentiation around putative centromeric regions. Mean pairwise F_ST_ estimated in 50 kb windows passing the minimum of 10 SNPs in the curated50 dataset and grouped by genomic region: annotated centromeric regions, 1 Mb pericentromeric flanks, and the remaining autosomal background. Negative window means were set to zero. There was no statistically detectable difference among the three distributions (Kruskal–Wallis test, P = 0.969).

### Supplementary Text S5. Runs of homozygosity

#### Runs of Homozygosity

ROH analyses were performed using the final filtered autosomal SNP dataset (1,324,912 SNPs). This dataset had previously been filtered to remove low-quality genotypes (DP < 5, GQ < 20), sites with more than 20% missing data, and variants with minor allele frequency below 2%. Two samples identified as contaminants during quality-control analyses were removed prior to ROH calling. All remaining individuals were retained irrespective of relatedness, as ROH analyses quantify individual genomic inbreeding and are therefore not biased by the inclusion of related samples. ROHs were identified using PLINK v1.9 [31] with the parameters described in the main text. Genomic inbreeding coefficients (FROH) were calculated as the proportion of the autosomal genome contained within ROHs.

**Table S8.**
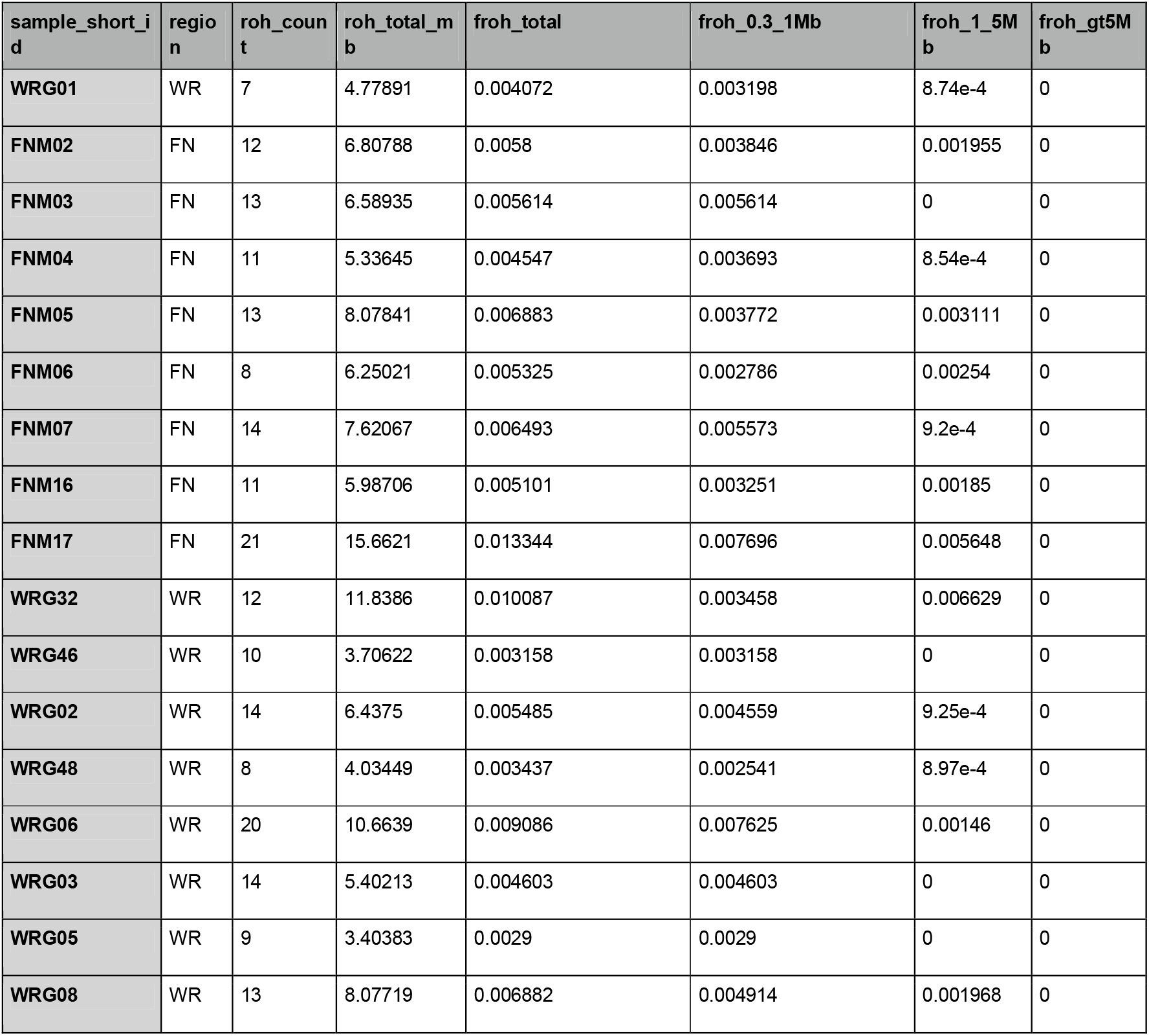
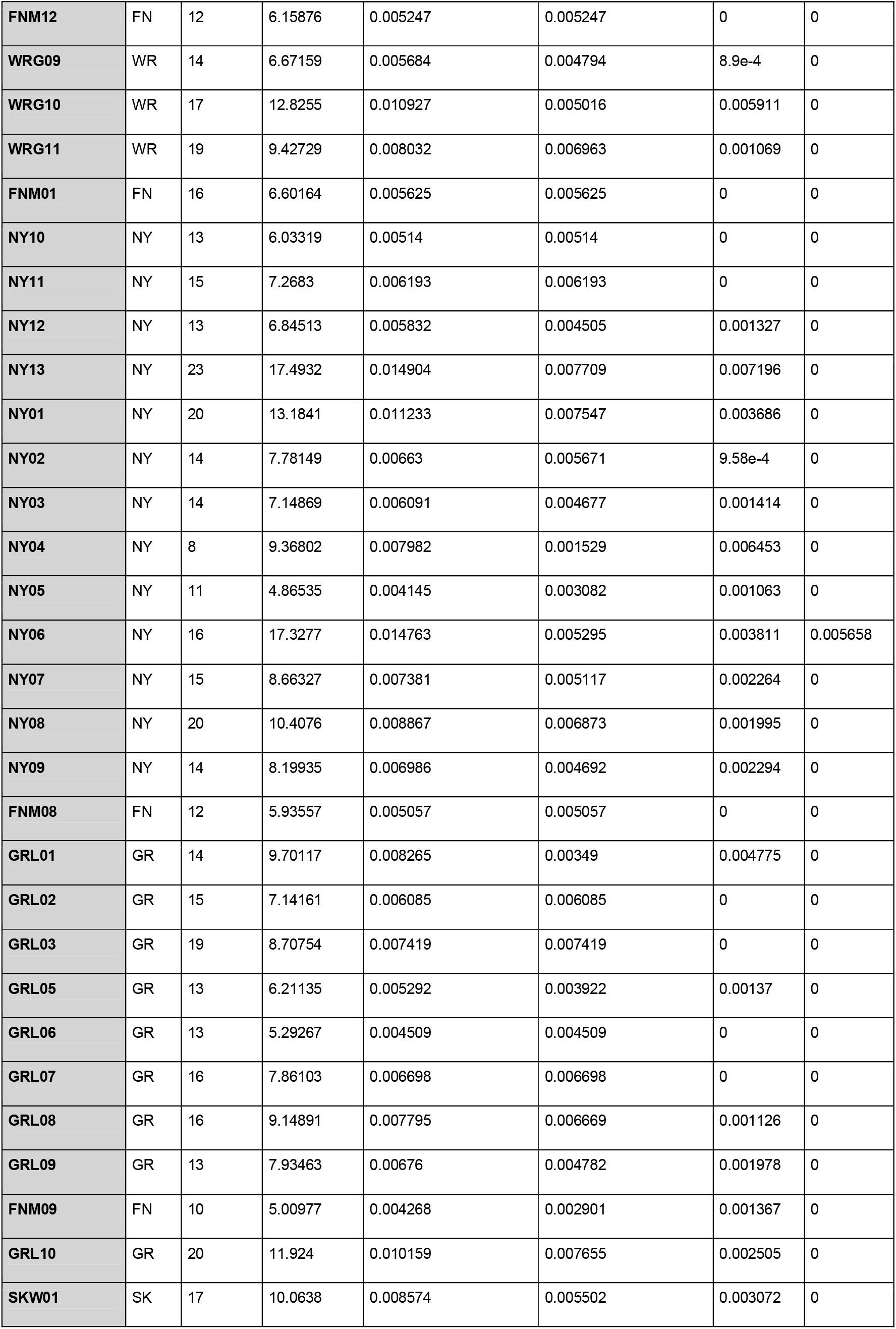
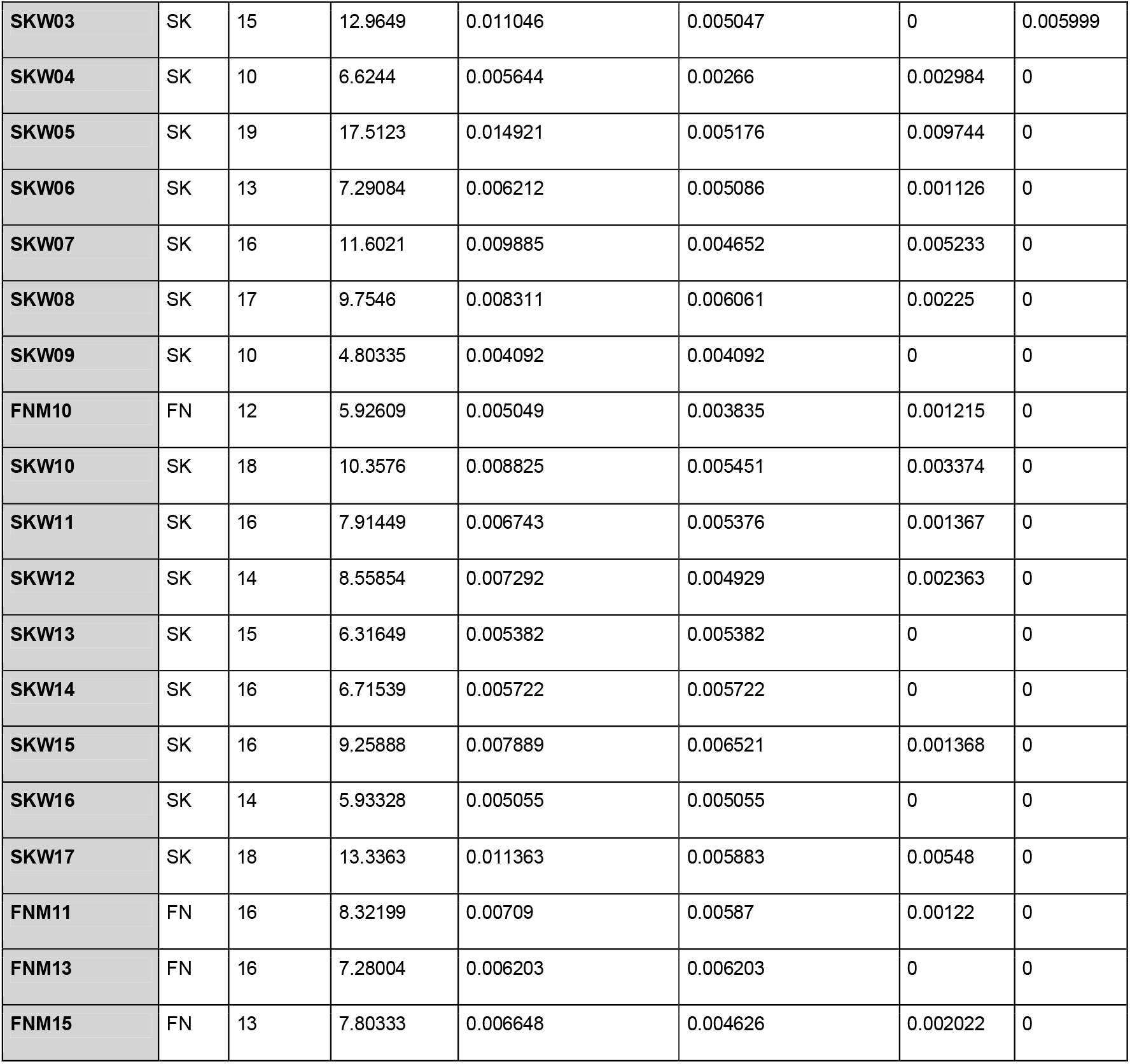
Individual runs of homozygosity summary. Individual-level ROH summaries, including number of ROH segments, total ROH length, genome-wide FROH, and FROH partitioned by ROH length class.

**Table S9.** Regional runs of homozygosity summary. Regional and all-sample summaries of ROH burden, including number of individuals, mean and median FROH, maximum FROH, mean and median ROH count, and mean and median total ROH length.

| region | n_individuals | mean_froh | median_froh | max_froh | mean_roh_count | median_roh_count | mean_roh_total_mb | median_roh_total_mb |
| --- | --- | --- | --- | --- | --- | --- | --- | --- |
| All | 66 | 0.007102 | 0.006561 | 0.014921 | 14.333333 | 14 | 8.335637 | 7.70108 |
| FN | 16 | 0.006143 | 0.005619 | 0.013344 | 13.125 | 12.5 | 7.210583 | 6.595495 |
| GR | 9 | 0.006998 | 0.00676 | 0.010159 | 15.444444 | 15 | 8.213657 | 7.93463 |
| NY | 13 | 0.008165 | 0.006986 | 0.014904 | 15.076923 | 14 | 9.583492 | 8.19935 |
| SK | 16 | 0.007935 | 0.00759 | 0.014921 | 15.25 | 16 | 9.312954 | 8.90871 |
| WR | 12 | 0.006196 | 0.005585 | 0.010927 | 13.083333 | 13.5 | 7.272263 | 6.554545 |

### Supplementary Text S6. GONE demographic inference

- Methods: recombination-rate estimation, chromosome subset, high-depth dataset, sensitivity runs

#### GONE analyses

Inference of recent effective population size using GONE requires an estimate of the recombination rate. Because no recombination map is currently available for snowy owl (*Bubo scandiacus*), recombination rates were approximated using chromosome-specific estimates from chicken (*Gallus gallus*) [40,41].

To minimize bias associated with differences in chromosome structure between species, we restricted analyses to a subset of snowy owl autosomes that satisfied four criteria:

1. One-to-one chromosomal homology with chicken based on whole-genome synteny analyses [23].
2. No evidence of chromosome fusion or fission relative to the homologous chicken chromosome.
3. Comparable chromosome morphology between species, with both chromosomes classified as acrocentric or subtelocentric [40].
4. Availability of published chromosome-specific recombination rate estimates for the homologous chicken chromosome.

Chromosome size and centromere position are major predictors of broad-scale recombination rate, partly because at least one crossover is generally required for proper meiotic segregation of each chromosome pair, and because crossover number and placement are constrained by chromosome length and arm structure [84]. This makes the recombination rate per Mb strongly dependent on chromosome size: small chromosomes tend to have high recombination rates per physical distance, whereas large chromosomes often contain broad regions of reduced recombination. In birds, microchromosomes therefore have substantially higher recombination rates per Mb than macrochromosomes and generally provide a more comparable chromosome class for cross-species approximation of recombination rates [85]. We treated acrocentric and subtelocentric chromosomes as comparable because both have terminal or subterminal centromere positions, whereas metacentric chromosomes were excluded when the homologous chromosome differed strongly in morphology. Macrochromosomes were excluded because recombination rates differ substantially between macrochromosomes and microchromosomes in birds [86,87], chromosome size and arm structure vary more among macrochromosomes, and only one macrochromosome pair met our synteny and morphology criteria. Chromosomes exhibiting evidence of fusion or fission [23], uncertain homology, strongly divergent morphology, or lacking published recombination-rate estimates were also excluded.

Application of these criteria resulted in retention of 17 snowy owl microchromosomes (11, 13–15, and 17–29) (Table S10). For each retained chromosome, the chromosome-specific recombination rate reported for the homologous chicken chromosome was assigned to the snowy owl chromosome. An arithmetic mean recombination rate was then calculated across all retained chromosomes, yielding an estimate of 6.63 cM/Mb. This value was used in all GONE analyses.

**Supplementary Table S10.** Chromosomes used for GONE demographic inference. snowy owl chromosomes retained for GONE analyses were selected based on one-to-one synteny with chicken, chromosome morphology, and availability of chromosome-specific recombination-rate estimates. Chromosome numbers in the snowy owl columns refer to the chromosome identifiers in the snowy owl reference assembly. Each retained snowy owl chromosome was assigned the recombination rate of its homologous chicken chromosome. The arithmetic mean recombination rate across the 17 retained chromosomes was 6.63 cM/Mb.

| snowy owl |  |  | Chicken |  |  |  |
| --- | --- | --- | --- | --- | --- | --- |
| Chromosome | Length Mb | Morphology | Homologous chromosome | Morphology | Length Mb | Recombination rate cM/Mb |
| 11 | 27.9 | Acrocentric | 12 | Acrocentric | 20.5 | 3.4 |
| 13 | 26.1 | Acrocentric | 10 | Acrocentric | 22.5 | 3.6 |
| 14 | 24.5 | Acrocentric | 11 | Subtelocentric | 21.9 | 2.9 |
| 15 | 22.4 | Acrocentric | 13 | Acrocentric | 18.9 | 3.3 |
| 17 | 20.1 | Acrocentric | 14 | Acrocentric | 15.8 | 4.3 |
| 18 | 19.1 | Acrocentric | 15 | Acrocentric | 13.0 | 4.1 |
| 19 | 18.3 | Acrocentric | 20 | Acrocentric | 13.9 | 4.0 |
| 20 | 17.6 | Acrocentric | 17 | Acrocentric | 11.2 | 4.6 |
| 21 | 14.6 | Acrocentric | 18 | Acrocentric | 10.9 | 4.7 |
| 22 | 13.5 | Acrocentric | 19 | Acrocentric | 9.9 | 5.3 |
| 23 | 12.1 | Acrocentric | 28 | Acrocentric | 4.5 | 11.9 |
| 24 | 11.6 | Acrocentric | 21 | Acrocentric | 6.9 | 8.2 |
| 25 | 10.0 | Acrocentric | 22 | Acrocentric | 3.9 | 14.3 |
| 26 | 9.5 | Acrocentric | 23 | Acrocentric | 6.0 | 8.7 |
| 27 | 8.7 | Acrocentric | 27 | Acrocentric | 4.7 | 10.8 |
| 28 | 8.5 | Acrocentric | 24 | Acrocentric | 6.4 | 8.3 |
| 29 | 8.0 | Acrocentric | 26 | Acrocentric | 5.1 | 10.3 |

The final GONE dataset contained 64 individuals after removal of two contaminated samples and two closely related individuals. SNPs were derived from the genotype-filtered autosomal dataset prior to minor allele frequency filtering, thereby maximizing the number of informative loci available for linkage disequilibrium analyses. No linkage pruning was performed, following recommendations for GONE analyses.

The primary GONE analysis was performed using 24 high-depth individuals. This dataset was selected after sensitivity analyses showed that analyses including all 64 individuals were sensitive to low-depth individuals and very rare variants. SNPs were derived from the genotype-filtered autosomal dataset, and no linkage pruning was performed, following recommendations for GONE analyses.

Sensitivity analyses were performed using both the full 64-individual dataset and the 24-individual high-depth dataset. For each dataset, analyses were run using polymorphic sites and using minor allele frequency thresholds of 0.005, 0.01, and 0.02. GONE inference was robust within the high-depth dataset across filtering thresholds, whereas the all-64 polymorphic-site and MAF >= 0.005 analyses inferred biologically implausible *N*_e_ values. In contrast, all-64 analyses with MAF >= 0.01 or MAF >= 0.02 produced trajectories broadly comparable to the high-depth analyses. We therefore used the 24-individual high-depth dataset as the primary demographic inference and retained the full sensitivity analysis as supporting information.

Long-term GONE trajectories extending beyond ~200 generations are shown for exploratory context only. Because GONE is based on linkage disequilibrium, inference becomes increasingly uncertain farther back in time as LD signals weaken and become more sensitive to recombination-rate assumptions, marker filtering, sample composition, and genotyping error. Older features were therefore not used as primary evidence for specific demographic events.

**Table S11.** GONE dataset and sensitivity summary. Summary of the primary GONE run and sensitivity analyses, including sample set, variant filter, number of inferred generations, generation time, present year, minimum and maximum Ne across the full output, most recent and oldest Ne estimates, maximum years before present, and most recent Ne shown in the plotted interval.

| run_id | sample_set | filter | n_generations | generation_time_years | present_year | min_ne_full_output | max_ne_full_output | most_recent_ne_full_output | oldest_ne_full_output | max_years_before_present_full_output | most_recent_ne_plotted |
| --- | --- | --- | --- | --- | --- | --- | --- | --- | --- | --- | --- |
| all64_poly | All 64 | Polymorphic sites | 791 | 8 | 2021 | 154.737 | 1208280 | 154.737 | 63164 | 6328 | 154.737 |
| all64_maf005 | All 64 | $MAF \geq 0.005$ | 791 | 8 | 2021 | 155.268 | 1545360 | 155.268 | 52013.6 | 6328 | 155.268 |
| all64_maf01 | All 64 | $MAF \geq 0.01$ | 791 | 8 | 2021 | 7964.47 | 29638.1 | 19831.7 | 29462 | 6328 | 19831.7 |
| all64_maf02 | All 64 | $MAF \geq 0.02$ | 755 | 8 | 2021 | 8818.94 | 32686.4 | 20042.9 | 32686.4 | 6040 | 20042.9 |
| highdepth24_poly | High-depth 24 | Polymorphic sites | 755 | 8 | 2021 | 8018.35 | 60170.5 | 18134 | 44122.3 | 6040 | 18134 |
| highdepth24_maf005 | High-depth 24 | MAF >= 0.005 | 755 | 8 | 2021 | 7623.54 | 69660.1 | 17157.1 | 47679.5 | 6040 | 17157.1 |
| highdepth24_maf01 | High-depth 24 | MAF >= 0.01 | 755 | 8 | 2021 | 7914.15 | 60659.4 | 16813.3 | 44849.7 | 6040 | 16813.3 |
| highdepth24_maf02 | High-depth 24 | MAF >= 0.02 | 755 | 8 | 2021 | 8694.29 | 57896 | 17401.4 | 41982.3 | 6040 | 17401.4 |

#### Data Availability and Online Data Tables

Extended tables underlying the analyses are provided as online data tables in the associated repository/archive. These include full sample metadata, genome-wide pixy window estimates, full pairwise KING relatedness estimates, ROH segment and density data, recurrent high-FST outlier annotations, and GONE sensitivity results. FASTQ provenance and ENA submission metadata are provided separately with the Zenodo data package (reserved 10.5281/zenodo.22825317).

Online Data Table 1. Full sample metadata and analysis inclusion table.

Online Data Table 2. Full pairwise KING kinship estimates.

Online Data Table 3. Full pairwise FST and dXY window table from pixy.

Online Data Table 4. Recurrent high-FST outlier windows and annotations.

Online Data Table 5. Full individual ROH segment table.

Online Data Table 6. ROH density by genomic window.

Online Data Table 7. GONE sensitivity-run estimates.

